# Calyculin A Induces Premature Chromosome Condensation and Chromatin Compaction in G_1_-Phase HeLa Cells without Histone H1 Phosphorylation

**DOI:** 10.1101/2025.07.16.665228

**Authors:** Natalia Y. Kochanova, Matthieu Vermeren, Bram Prevo, Ipek Ustun, Shaun Webb, Linfeng Xie, William C. Earnshaw, James R. Paulson

## Abstract

We show here that treatment of HeLa cells with calyculin A, an inhibitor of Protein Phosphatases 1 and 2A, induces premature chromosome condensation (PCC) at any point in interphase of the cell cycle. Chromosomes in G_1_-phase PCC closely resemble metaphase chromatids in the light microscope, and measurements using FLIM-FRET show that they have the same level of chromatin compaction as metaphase chromosomes. However, histone H1 is not phosphorylated in G_1_- or early S-phase PCC. These results suggest that H1 phosphorylation is not required for mitotic chromosome condensation and chromatin compaction. They also confirm that Cdk1/cyclin B, which directly phosphorylates histone H1, is not active in G_1_ and thus is not essential for G_1_- PCC. We suggest that induction of G_1_-PCC involves protein kinases or other factors that are either held in an inactive state by protein phosphatases, or constitutively active but countered by phosphatases. The same factors may be involved in the onset of normal mitosis, becoming active when protein phosphatases are downregulated. Induction of PCC with calyculin A should provide a useful system for identifying and studying the biochemical pathways that are required for mitotic chromosome compaction, nuclear envelope breakdown, and other events of mitosis.

## Introduction

The onset of mitosis is initiated by activation of Cdk1/Cyclin B kinase (Norbury and Nurse, 1992), which leads to the phosphorylation of many proteins either by Cdk1 itself or by secondary protein kinases (Gurley et al., 1978; Dorée and Galas, 1994; Stukenberg et al., 1997; Ferrari, 2006; Malik et al., 2008; Hegemann et al., 2011). This protein phosphorylation triggers the characteristic events of mitosis including nuclear envelope breakdown (Ward and Kirschner, 1990; Heald and McKeon, 1990; Peter et al., 1990), nuclear pore complex disassembly (Laurell et al., 2011), chromosome condensation (Wei et al., 1998, 1999; Van Hooser et al., 1998), cessation of transcription (Gottesfeld and Forbes, 1997), reorganization of the Golgi (Shorter and Warren, 2002) and reorganization of the cytoskeleton (Chou et al., 1989; Yamaguchi et al., 2005). At the end of mitosis, Protein Phosphatase 1 (PP1) and other protein phosphatases dephosphorylate these proteins, and the cell returns to interphase (e.g., Glass and Gerace, 1990; Foisner and Gerace, 1993; Paulson et al., 1996; Thompson et al., 1997; Nelson et al., 1997; de Castro et al., 2016, 2017; Huguet et al., 2019).

Histones are the most abundant proteins in eukaryotic chromatin. DNA is wrapped around core histone octamers to form nucleosomes (e.g., Luger et al., 1997; Kornberg and Lorch, 1999) that are arranged along the DNA in a “string of beads” or 11 nm fiber. Evidence for higher-order chromatin structure, requiring histone H1 (Thoma et al., 1979), comes from electron microscopy and small-angle x-ray scattering (e.g., DuPraw, 1965, 1970; Marsden and Laemmli, 1979; Rattner and Hamkalo, 1979; Langmore and Paulson, 1983; Paulson and Langmore, 1983; Robinson et al., 2008).

Ricci et al. (2015) report that the 11 nm fiber is compacted into heterogeneous “clutches” of nucleosomes. Recent cryo-EM studies suggest that in highly condensed metaphase chromosomes, nucleosomes form an amorphous “polymer melt” (e.g., Eltsov et al., 2008; Maeshima et al., 2010; Joti et al., 2012). This has been suggested to involve phase-separation of the nucleosomes (Gibson et al., 2019; Schneider et al., 2022).

During mitosis, several histones are phosphorylated. Histone H2A is phosphorylated at T120 by Bub1 kinase. This helps to recruit Topoisomerase 2α and the chromosomal passenger complex (CPC) to centromeres (Akera and Watanabe, 2016; Abad et al., 2019; Zhang et al., 2020). Histone H3 is phosphorylated at T3 by Haspin kinase (Dai et al. 2005). This helps recruit the CPC as well as Shugoshin (Kawashima et al., 2010; Yamagishi et al., 2010). H3 is also phosphorylated at Ser10 and Ser28 by Aurora B kinase (Hsu et al., 2000). Histone H1 is mainly phosphorylated by Cdk1/cyclin B (Garcia et al., 2004; Deterding et al., 2008), which has long been known as the mitotic histone H1 kinase (e.g., Lake and Salzman, 1972; Labbé et al., 1989; Langan et al., 1989). Indeed, H1 kinase activity was a key assay used in the initial purification of Cdk1/cyclin B (Lohka et al., 1988; Gautier et al., 1988; Gautier et al., 1990). H1 is rapidly dephosphorylated at the end of mitosis, mainly by Protein Phosphatase 1 in HeLa cells (Paulson et al., 1996).

Many studies have investigated the premature chromosome condensation (PCC), that occurs when a metaphase-arrested cell is fused with an interphase cell (Johnson and Rao, 1970), thereby introducing active Cdk1/cyclin B into the latter (Rao, 1990). PCC can also be induced by treating interphase cells with calyculin A, an inhibitor of Protein Phosphatases 1 and 2A (Ishihara et al., 1989; Gotoh et al., 1995; Alsbeih and Raaphorst, 1999; Kanda et al., 1999). Calyculin A-induced PCC has been used for cytogenetic analysis and studies of chromosome aberrations (e.g., Suzuki et al., 2001; Bezrookove et al., 2003; Srebniak et al., 2005; Miura and Blakely, 2011), to investigate radiation sensitivities (e.g., Coco-Martin and Begg, 1997; Pathak et al., 2009), and to study chromosome dynamics during S-phase (Gotoh, 2007). The protein phosphatase inhibitors okadaic acid and cantharidin can also induce PCC (e.g., Gowdy et al., 1998; Prasanna et al., 2000).

In this paper, we have used FLIM-FRET (Fluorescence Lifetime Imaging Microscopy-Förster Resonance Energy Transfer) (Llères et al., 2007) to measure chromatin compaction in live cells. Our protocol employs histone H2B molecules fused to donor and acceptor fluorescent proteins at their C- and N- termini, respectively. In FRET, photons from the donor excite the acceptor, which must be within 10nm.

Measurement of FRET efficiency indicates how close donor and acceptor proteins are to one another in adjacent nucleosomes. FRET does not occur between donor and acceptor in the same nucleosome because their spatial separation is too great. Determination of nucleosome proximity in this way has previously been used to quantify mitotic chromosome compaction at different stages of cell division, and to differentiate between eu- and heterochromatic regions in interphase cells (Llères et al., 2009; DuPont et al., 2023).

Here, we have explored the phenomenon of calyculin A-induced PCC. We find that calyculin A induces PCC at all stages of the cell cycle, but histone H1 is not phosphorylated in G_1_- and early S-phase PCC. Remarkably, light microscopy and FLIM-FRET measurements reveal no difference in chromosome condensation or chromatin compaction between G_1_-PCC and metaphase chromosomes even though H1 is not phosphorylated.

## Materials and Methods

### Chemicals and Media

For treatment of H-HeLa or HeLa S3, calyculin A was prepared as a 100 μM solution in methanol, stored at 2°C, and used within 2 weeks. For treatment of HeLa CDK1^as^ cells, calyculin A was dissolved at 100 μM in dimethyl sulfoxide (DMSO) and stored at -80°C. For a 5 M stock solution of sodium butyrate, butyric acid was dissolved in 0.9% NaCl and 20 mM sodium phosphate, adjusted to pH 7.4 with NaOH, and stored at room temperature. Thymidine (TdR) was dissolved to 100 mM in 0.9% NaCl, filter sterilized and stored at 2°C or prepared as a 100 mM or 24 mM solution in water, filter sterilized and stored at −20°C. Nocodazole stock solutions were either 5 mg/mL or 1 mg/mL in DMSO and stored at −20°C. Roscovitine (Meijer et al., 1997) was prepared as a 50 mM stock solution in DMSO and stored at −20°C. 1NM-PP1 (Bishop et al., 2000) was synthesized as described previously (Keaton et al., 2023), dissolved in DMSO and stored at -20°C. Deoxycytidine (CdR) was dissolved in water as a 24 mM stock solution and stored at −20°C. Blasticidin was prepared as a 25 mg/mL solution in water and stored at −20°C. Media and components were obtained from Gibco/Invitrogen or Sigma. All other reagents were obtained from Sigma unless otherwise noted.

### Cell Culture and Synchronization

The suspension HeLa cultures used were either H-HeLa (Medappa et al., 1971) or HeLa S3. The adherent cell line used was HeLa CDK1^as^ (Rata et al. 2018). H-HeLa cells were grown in Eagle’s MEM as previously described (Paulson et al., 1996). HeLa S3 cells were grown in RPMI-1640 supplemented with penicillin/streptomycin and 10% fetal bovine serum and diluted daily to 2.0 – 2.5 × 10^5^/mL. Monolayers of HeLa CDK1^as^ were cultured in DMEM supplemented with 10% FBS and penicillin/streptomycin and split every 2-3 days.

For synchronization of suspension cultures in G_1_-phase, cells were treated with 5 mM sodium butyrate for 20 hrs (e.g., Darzynkiewicz et al., 1981; Wintersberger et al., 1983; Han et al., 1987; Vaziri et al., 1998; Chen and Faller, 2005). Rapid and efficient release from the butyrate block was achieved by pelleting the cells, washing twice with 0.9% NaCl, and resuspending the cells in fresh medium. Cells were synchronized in S-phase by treatment with 2.5 mM thymidine for 20-24 hrs (Xeros, 1962). For metaphase arrest, cells were first blocked with thymidine, then released and arrested with nocodazole as described previously (Paulson, 1980). Mitotic indices for H-HeLa and HeLa S3 were determined as previously described (Paulson, 2007) and were typically 80- 95% and 95-98%, respectively.

To obtain G_1_ or G_1_-PCC cultures of adherent HeLa CDK1^as^ cells, a 2.5 mM thymidine block was implemented for 18 hrs, after which the thymidine was washed out and 24 μM deoxycytidine and 24 μM thymidine (Heintz et al., 1983) were added together with 2 μM 1NM-PP1. After 8 hrs, the culture flasks were shaken and then washed 3 times with 50% conditioned media (consecutive washes of 2, 5 and 3 min in the incubator) to get rid of dead cells and remove 1NM-PP1 so that the cells could progress into mitosis. After the third wash, 50% conditioned media was added. 1 hr later, mitotic cells were shaken off, 2.5 mM thymidine was added, and the cells allowed to re-adhere in a new flask. After an additional 3 hrs, the adhered cells were in G_1_. Non-adhered cells were removed with the medium and fresh medium (containing thymidine) was added.

Optionally, calyculin A was also added to these G_1_ cells for 2 hrs.

For each replicate of the FLIM-FRET experiment, the five cell lines (four containing fusion proteins plus the parent HeLa CDK1^as^) were treated as described, at 45- minute intervals, after which cells were imaged in 50% conditioned Liebovitz L-15 medium without phenol red, supplemented with 10% FBS and penicillin/streptomycin.

To block an adherent HeLa CDK1^as^ culture in metaphase, 0.25 µg/mL nocodazole was added to a non-confluent culture for 14 hrs. For metaphase spreads, mitotic cells were obtained by shake-off.

### Establishing cell lines expressing histone H2B and fluorescent proteins

mNeonGreen and mScarlet-I proteins were chosen for FRET imaging (McCullock et al., 2020) and several fusion proteins were constructed: H2B-mNeonGreen-T2A- mScarlet-I-H2B; H2B-mNeonGreen-T2A-mScarlet-I; H2B-mNeonGreen; and mNeonGreen-linker-mScarlet-I (Llères et al., 2007, 2009) with 7 amino acid linker GGGGSGG. These constructs were cloned into the pMK457 piggyback transposase vector (a gift from Masato T. Kanemaki), restricted with EcoRI and BamH1, under an EF1 promoter. Primers for cloning into the pcDNA5 FRT vector (Thermo Fisher Scientific), into which the constructs were cloned first before subsequent re-cloning into pMK457, are listed in Table S1. The mScarlet-I-containing plasmid was a gift from the lab of Alistair McCormick, and the mNeonGreen-containing plasmid was a gift from the lab of Julie Welburn. The H2B-containing plasmid was a gift from Patrick Heun. The cloning was done with the InFusion system (Takara Bio) according to the manufacture’s protocols.

For transfection, HeLa CDK1^as^ cells were seeded at 1.92 x 10^5^ cells per well into 6 well plates and transfected the next day with FuGENE® 6 transfection reagent according to the manufacturer’s protocol. We used 3 µg of plasmid (0.2-1 mg/mL concentration, A260/A280 ratio 1.7-1.9) and FuGENE® 6 : DNA in a 3:1 ratio. After two days, cells from each well were harvested in 2 mL of medium containing trypsin and partially transferred to 10 cm dishes at three different dilutions: 1:40 (250 μL,) 1:20 (500 μL), and 1:10 (1000 μL). Medium was then added to give a total volume of 10 mL in each dish. Blasticidin was added the next day at a concentration of 5 μg/mL and the medium was changed every 3-4 days when dead cells accumulated. For each cell line, when the colonies reached sufficient size, between 5 and 40 colonies were picked with tips, grown separately, and screened by fluorescence. Chosen clones were grown further with 5 μg/mL blasticidin.

### Western blotting

For each of the four cell lines constructed, and for the control parental cell line, 5 x 10^6^ cells were harvested and frozen. The pellets were resuspended in 300 μL of lysis buffer consisting of 20 mM Tris (pH 7.4), 0.1% Triton X-100, 150 mM NaCl, 1 mM EDTA (Thermo Fisher Scientific), 1 μg/mL protease inhibitor cocktail (chymostatin, leupeptin hemisulfate, antipain dihydrochloride, pepstatin A), and 0.15 mM PMSF (Harlow and Lane, 1999). The samples were sonicated for 15 cycles of 30 sec ON/30 sec OFF (“high” mode) in a Bioruptor® Plus (Diagenode) and centrifuged for 5 min (2300 x g, 4°C). 4x LDS sample buffer (Invitrogen), was added to the supernatant to give a final concentration of 1x LDS, supplemented with 0.1 M DTT (Melford). Samples were boiled for 5 min at 95°C and loaded on precast 4-12% BisTris gels (Invitrogen) together with SeeBlue Plus2 Pre-stained Protein Standards (Thermo Fisher Scientific) as markers. After electrophoretic separation, the proteins were transferred to Amersham Protran 0.45 NC nitrocellulose membranes (2 hrs, 100 V). The membranes were blocked with 5% milk. Primary antibodies in 1% milk were applied overnight: rabbit anti-H2B (Abcam ab1790, 0.1 μg/mL); rabbit anti-mNeonGreen (Cell Signalling, 53061, 1:1000); mouse anti-RFP (ChromoTek, 6G6, 1:1000); or rabbit anti-nucleolin (Abcam ab22758, 1 μg/mL). The next day the membranes were washed three times in PBS containing 0.2% Tween-20 (PBST) and incubated for 1 hr with secondary antibodies in 1% milk: goat anti-rabbit (Alexa 680 or 800, A32734 and A32735, Invitrogen); or donkey anti-mouse (680RD, 926-68072, IRDye®). Membranes were finally washed twice in PBST and imaged on the LiCor Odyssey CLx Infrared Imaging System (LICORbio^TM^). Images were processed with Image Studio™ Software (LICORbio^TM^).

### Histone Extraction and Analysis of Histone H1 Phosphorylation

For analysis of histone H1 phosphorylation, crude metaphase chromosomes, prematurely condensed chromosomes, and/or interphase nuclei were prepared by the procedure for isolating chromosome clusters (Paulson, 1982; Paulson et al., 1996). Lysis solutions contained 2 mM p-chloromercuriphenyl sulfonate (PCMPS) or 2 mM p- hydroxymercuribenzoate (PHMB) to block histone dephosphorylation (Paulson, 1980). Histones were extracted from pelleted chromosomes or nuclei with 0.2 M H_2_SO_4_ as previously described (Paulson, 1980). Samples for electrophoresis were freeze-dried in 1.5 mL microcentrifuge tubes using a CentriVap (Labconco) and redissolved in 3-4 μL of sample buffer. Gels for acid-urea electrophoresis (Panyim and Chalkley, 1969) were 16 cm long. They contained 15% acrylamide, 0.1% N,N’-methylene-bis-acrylamide, 2.5 M urea and 5.4% acetic acid, and were pre-electrophoresed (before loading samples) as described by Paulson and Higley (1999). Samples were then loaded and the gels run with fresh electrode well buffer at 300 V for 5-6 hrs until the blue component of the methyl green marker reached the bottom of the gel. All gels were stained with 0.1% Coomassie Brilliant Blue R250 (BioRad) in 50% methanol/10% acetic acid and destained in 5% methanol/10% acetic acid.

Note that on acid-urea gels the ratio of phosphorylated and unphosphorylated histone H1 can be clearly seen by a mobility shift. These gels have the advantage that they can reveal when H1 is fully phosphorylated. Absence of unphosphorylated H1 would be difficult to show using ^32^P-labelling or western blotting with anti-phospho-H1 antibodies.

### Microscopy of Fixed Cells and Chromosome Spreads

For microphotography, cells in a culture aliquot of at least 5 mL were pelleted and resuspended in 2 mL 75 mM KCl. After incubation for 15 min at 37°C they were fixed by addition of 200 µL of fresh fixative (3:1 (v/v) methanol:acetic acid). Fixed cells were then pelleted and gently resuspended in fresh fixative three more times. Droplets of the final fixed cell suspension were dropped from a height of 20 cm onto cold, wet microscope slides to spread the chromosomes (Musio et al., 1997). Slides were air dried, mounted and stained with VectaShield containing DAPI (VectorLabs), and viewed in a DeltaVision microscope with Olympus UPlanXApo 100X oil objective, NA 1.45.

For immunofluorescence microscopy of chromosomes and nuclei (from G_1_, G_1_- PCC or metaphase cultures), fixed cells were analogously dropped on cold wet coverslips which were then dried and put in PBS. Spreads were blocked with 5% BSA at 37°C, followed by incubation with anticentromere antibodies (ACA) (1:500 dilution) (Earnshaw and Rothfield, 1985). Coverslips were washed 3 times with PBS and incubated with secondary goat anti-human IgG Alexa Fluor^TM^ 488 antibodies (A11013, Invitrogen). Following 2 additional washes with PBS, samples were mounted in Vectashield with DAPI and imaged on the DeltaVision microscope. Images were subjected to deconvolution and maximum intensity projections were generated in ImageJ. For statistics, the spreads or cells were viewed in both channels (DAPI and FITC) and classified under the microscope.

For fluorescence microscopy of intact cells, the cells were rinsed with PBS, fixed with 4% formaldehyde for 8 minutes at 37°C, rinsed with PBS twice and mounted in Vectashield with DAPI. A single plane was imaged on the DeltaVision microscope with Olympus UPlanXApo 60X oil objective, NA 1.42. Images were processed in ImageJ.

### FLIM-FRET

For FLIM-FRET, cells were imaged using a Leica Stellaris FALCON confocal microscope with a 63x oil immersion objective (63x HC PL Apo CS2/1.40), equipped with a tunable white light laser and FLIM-designed hybrid detectors. Imaging was performed in a black-walled incubation chamber at 37°C with humidified atmosphere and constant 5% CO_2_ supply. Samples were imaged at a laser frequency of 20 mHz, and grabbed in photocounting mode ensuring no pile-up. Pulse intervals were 50 ns, which is ideal for lifetimes ranging between 0 and 15 ns. Each 16-bit image was scanned at 3 line and 3 frame repetitions at a speed of 100 Hz, to ensure statistically useful photon counts. All images acquired were 256 x 256 pixels. Each pixel was 724 x 724 nm in size, ensuring Nyquist sampling for whole nuclei (diameter size around 20 μm) and chromosome clusters (thickness size around 10 μm). For all cell lines, only the mNeonGreen channel was imaged, except for those cells expressing mNeonGreen-linker-mScarlet-I and the wild-type HeLa CDK1^as^ cells. In those cases, both mNeonGreen and brightfield channels were imaged to distinguish between interphase, mitotic, and G_1_-PCC cells.

For each replicate of the FLIM experiment, for each of the five cell lines, 20 images were acquired for each condition (G_1_, G_1_-PCC, and nocodazole-arrested). Nuclei or chromosome clusters for the H2B-mNeonGreen/mScarlet-I-H2B, H2B- mNeonGreen/mScarlet-I, and H2B-mNeonGreen-expressing cell lines, or whole cells for the mNeonGreen-linker-mScarlet-I-expressing cell line, were segmented. Lifetimes were calculated for each image as a mono-component exponential decay fit using Leica’s LASX small molecule software (Alvarez et al., 2019). Lifetime images were exported from this software and processed in ImageJ. Example lifetime distributions were exported from the software as well, normalized to maximum and plotted in R. The center of mass for each distribution was calculated and plotted as well. For FRET ratio calculations, mean lifetimes for each replicate were used. FRET efficiency was calculated as 1 − (τ_DA_/τ_D_) where τ_DA_ is the fluorescence lifetime of the donor (mNeonGreen) in the presence of acceptor (mScarlet-I) and τ_D_ is the fluorescence lifetime of the donor in the absence of acceptor (Llères et al., 2009; Prevo and Peterman, 2014).

Statistical analysis was performed and data was plotted in R, using the ggplot2 package (R Core Team, 2025).

## Results

### Calyculin A induces PCC in interphase HeLa cells

To test the ability of calyculin A to induce premature chromosome condensation (PCC) in HeLa cells, an exponentially growing, unsynchronized suspension culture was treated with 200 nM calyculin A and sampled at various times. This treatment clearly induced PCC (Fig. 1A). Nuclear envelope disassembly and condensed chromosomes were observed in virtually all cells within 3 hrs (Fig. 1B). Fig. 1C shows that ≥20 nM calyculin A is sufficient to induce PCC in nearly all cells within 3 hrs.

**Fig. 1.**
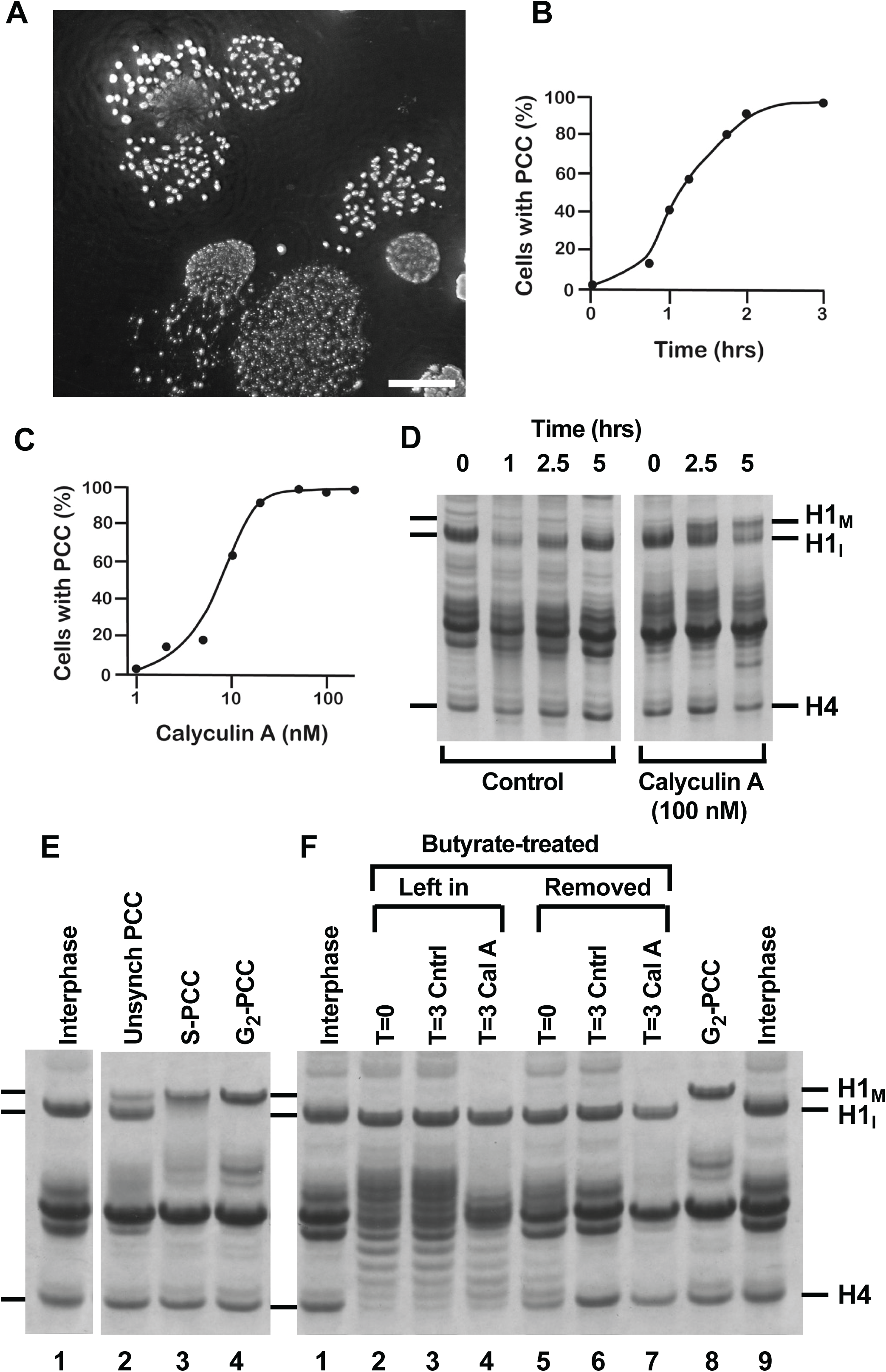
Induction of premature chromosome condensation (PCC) with calyculin A and analysis of histone H1 phosphorylation. (A) Micrograph of unsynchronized HeLa cells, spread and stained with DAPI, after 3 hrs treatment with 100 nM calyculin A. Some cells show condensed chromosomes (G_1_- and G_2_-PCC) and others show “pulverized chromosomes” (S-PCC). No interphase nuclei are seen. Scale bar represents 25 μm. (B) Percentage of unsynchronized HeLa cells displaying PCC as a function of the time of treatment with 200 nM calyculin A. (C) Percentage of unsynchronized cells displaying PCC after 3 hrs treatment with various concentrations of calyculin A. (D) Acid-urea gel electrophoresis of histones extracted from unsynchronized cells treated with 0.25 µg/mL nocodazole alone (control, left panel) or with both nocodazole and 200 nM calyculin A (right panel). H1_M_ and H1_I_ indicate the positions of mitotic (phosphorylated) and interphase histone H1, respectively. After 5 hrs with calyculin A, nearly 100% of cells exhibit PCC, but only some contain H1_M_. In this and subsequent figures, only the gel portions containing the histones are shown. (E), (F) Acid-urea gel electrophoresis of histones in PCC induced by treatment with 100 nM calyculin A for 3 hrs. (E) Histone H1 is unphosphorylated in interphase cells (lane 1). H1 phosphorylation is only partial in PCC induced in unsynchronized cells (lane 2), nearly complete in S-phase PCC (lane 3), and complete in G_2_-phase PCC (lane 4). (F) Histone H1 is not phosphorylated in G_1_- phase PCC. Cells were blocked in G_1_-phase by treatment with 5 mM sodium butyrate for 20 hrs, after which the butyrate was either left in (lanes 2-4) or removed by pelleting and washing the cells (lanes 5-7) before starting the calyculin A treatment. Histones were extracted at the start (T=0), after further incubation of the cells without treatment for 3 hrs (T=3 Ctrl), or after treatment for 3 hrs with 100 nM calyculin A (T=3 CalA).

In a separate experiment (Fig. 1D), cells were incubated either without (control) or with 200 nM calyculin A. At various times histones were extracted and analyzed by acid-urea polyacrylamide gel electrophoresis (Panyim and Chalkley, 1969; Paulson and Higley, 1999). Phosphorylated histone H1 (H1_M_) is substantially more abundant in the presence of 200 nM calyculin A than in its absence, but even after 5 hrs treatment, a significant amount of unphosphorylated H1_I_ remains (Fig. 1D, rightmost lane).

### Histone H1 phosphorylation in PCC depends on the cell cycle phase

The observation that histone H1 is not completely phosphorylated (Fig. 1D) when PCC is induced in virtually all cells of an unsynchronized HeLa culture suggested that H1 phosphorylation in PCC might be cell cycle dependent.

To test this possibility, HeLa cultures were synchronized with respect to the cell cycle, treated with calyculin A for 3 hrs, and their histones extracted and analyzed on acid-urea gels. S-phase cells were obtained by treatment with 2.5 mM thymidine, which suppresses cytidine (and therefore DNA) synthesis and blocks cells in S-phase (Xeros, 1962). G_2_-phase cells were obtained by release from the thymidine block (Fig. 1E).

As previously seen in Fig. 1D, unsynchronized interphase cells not treated with calyculin A lack histone H1 phosphorylation (Fig. 1E, lane 1), whereas those treated with calyculin A contain both phosphorylated and unphosphorylated H1 (H1_M_ and H1_I_, respectively) (Fig. 1E, lane 2). By contrast, calyculin A treatment of cells in S-phase (Fig. 1E, lane 3) or G_2_ (lane 4) leads to complete or nearly complete histone H1 phosphorylation (H1_M_).

These results suggested that it might be G_1_-PCC that lack phosphorylated histone H1 when an unsynchronized culture is treated with calyculin A (e.g., Fig. 1D and Fig. 1E, lane 2). To test this, HeLa cells were synchronized in G_1_-phase by treating a log-phase culture with 5 mM sodium butyrate for 20 hrs (cf. Darzynkiewicz et al., 1981; Wintersberger et al., 1983; Han et al., 1987; Vaziri et al., 1998; Chen and Faller, 2005).

Butyrate was left in half of this culture but removed from the other half by pelleting, washing with PBS, and resuspending the cells in fresh medium. For each half, three samples were prepared. Histones were extracted from one aliquot at the start, from a second aliquot after further incubation for 3 hrs without treatment, and from a third aliquot after treatment for 3 hrs with calyculin A. Analysis of these samples on acid urea gels is shown in Fig. 1F, where the three types of samples are labeled T=0, T=3 Ctrl, and T=3 Cal A, respectively. Lanes 2 – 4 show the results for cells that remained in 5 mM butyrate and lanes 5 – 7 show the results when butyrate was removed. In neither case is histone H1 phosphorylated following calyculin A treatment (Fig. 1F, lanes 4 and 7). For comparison, histones from interphase cells (with H1_I_) are shown in lanes 1 and 9, and histones from G_2_-phase PCC (with H1_M_) are shown in lane 8.

### In PCC, histone H1 is not phosphorylated until mid S-phase

It has been reported that sodium butyrate arrests HeLa cells in G_1_-phase before the restriction point, about 4-6 hrs before the start of S-phase (Han et al., 1987; Gong et al., 1994). The results in Fig. 1E and F suggest that only at some time after that restriction point is calyculin A treatment able to induce histone H1 phosphorylation.

To elucidate when in the cell cycle H1 can become phosphorylated in PCC, cells were synchronized in G_1_ by treatment with butyrate for 20 hrs. Butyrate was then removed, and half the culture was treated with thymidine, which blocks cells in S-phase, while the other half was treated with nocodazole, which blocks cells in mitosis. Samples were taken for determination of mitotic index at intervals of 4 hrs. At the same time points, aliquots were treated with 100 nM calyculin A for 4 hrs, and observations of S- PCC were used to determine the percentage of cells in S-phase. S-PCC is recognizable by the presence of so-called “pulverized chromosomes” (Johnson and Rao, 1970; Rao, 1990). The remainder of each calyculin A-treated aliquot was acid-extracted for analysis of H1 phosphorylation on acid-urea gels.

Fig. 2A shows the percentage of cells in S-PCC as a function of time for the thymidine-treated culture (●─●) and the nocodazole-treated culture (▴─▴). In both, it begins to rise at about T=4 hrs, where T=0 hrs is the time when butyrate was removed. The proportion of cells undergoing S-PCC is 50% at T=8 hrs after thymidine washout and reaches a maximum (about 85%) at T=16 hrs. After T=20 hrs, the percentage of S- PCC remains high in the thymidine-treated culture because the cells are blocked in S-phase, but in the nocodazole-treated culture it has begun to fall off as cells complete S-phase and enter G_2_.

**Fig. 2.**
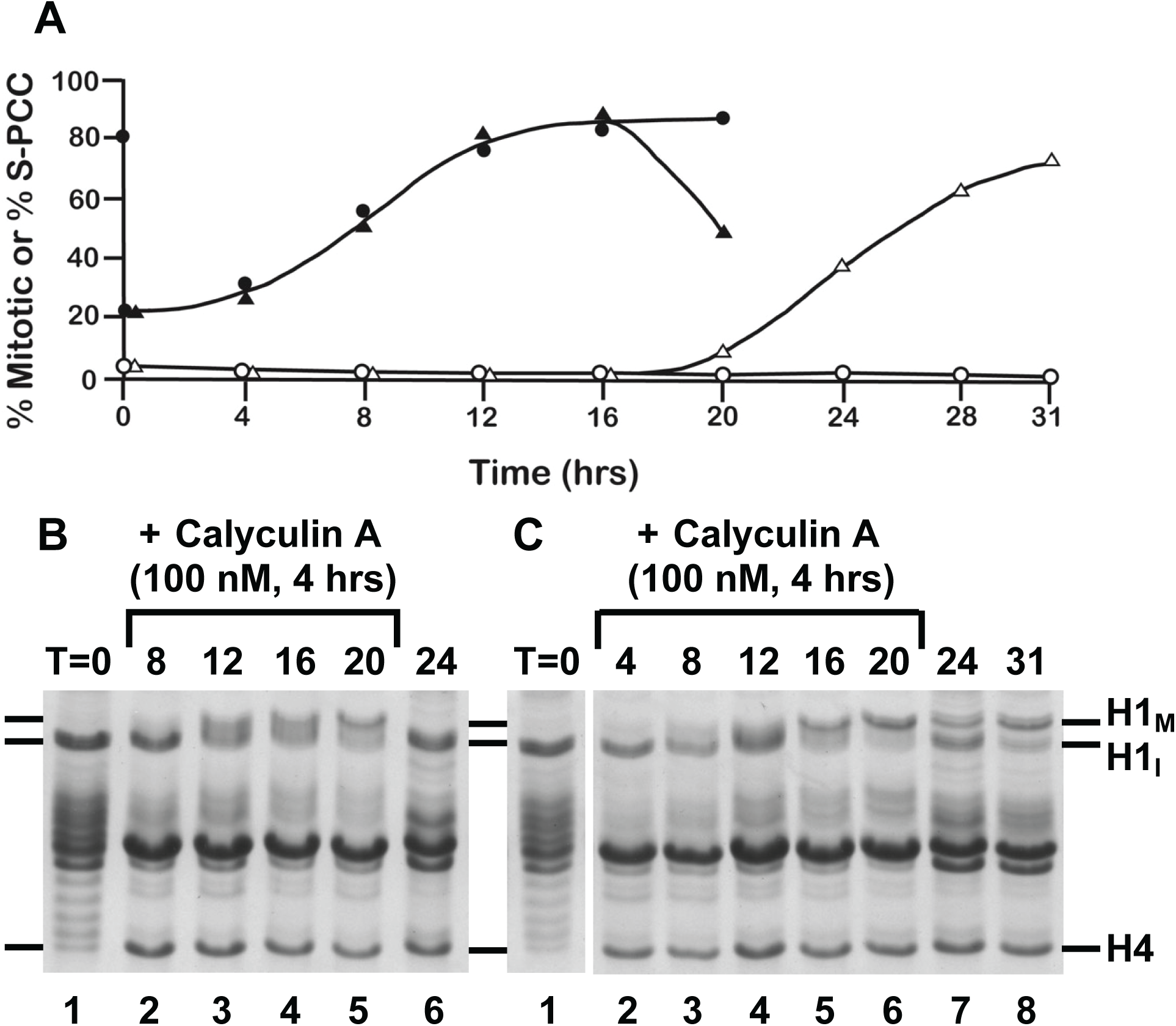
Histone H1 phosphorylation is not observed in HeLa PCC until the cells have progressed at least four hours into S-phase. (A) Cells synchronized in G_1_-phase by butyrate treatment and then released at T=0 hrs were treated with either 2.5 mM thymidine or 0.25 µg/mL nocodazole. At intervals of 4 hrs mitotic indices were determined: nocodazole-treated culture (Δ─Δ); thymidine-treated culture (○─○). Also at 4 hr intervals, aliquots were treated for 4 hrs with 100 nM calyculin A, and the percentage of cells in S-phase was determined as the percentage displaying “pulverized chromosomes”: nocodazole-treated culture (▴─▴); thymidine-treated culture (●─●). (B) Analysis of histones from the thymidine-treated culture, taken at the times shown. (C) Analysis of histones from the nocodazole-treated culture, taken at the times shown. Gel samples from T=4 to T=20 hrs are from culture aliquots treated 4 hrs with calyculin A, and for these T indicates the time (hrs) at which the calyculin A treatment was started. Samples labeled T=0, T=24 and T=31 were taken at those times (hrs) from cells that were not treated with calyculin A. Histones were extracted from chromosomes or nuclei with 0.2 M H_2_SO_4_ and separated on acid-urea gels.

Fig. 2A also shows the percentage of cells arrested in mitosis for the thymidine-treated culture (○─○) and the nocodazole-treated culture (Δ─Δ). The nocodazole-treated culture begins to enter mitosis at T=20 hrs and reaches a mitotic index of 71% at T=31 hrs. The thymidine-treated cells never reach mitosis because they are blocked in S-phase.

Figs. 2B and C show the extent of histone H1 phosphorylation in the various samples. Without calyculin A treatment, H1 never becomes phosphorylated in the thymidine-treated culture (Fig. 2B, lane 6) because the cells are blocked in S-phase. In the nocodazole-treated culture (without calyculin A), some H1 is phosphorylated at T=24 hrs (Fig. 2C, lane 7) and the majority is phosphorylated at T=31 hrs (Fig. 2C, lane 8) because a significant percentage of the cells have entered mitosis and have been arrested there by the nocodazole.

In the calyculin A-treated samples, T indicates the time at which the 4 hr calyculin A treatment was *started*. Little or no H1 phosphorylation is seen at T=8 hrs in either culture (Fig. 2B, lane 2; Fig. 2C, lane 3), even though about 50% of the cells have reached S-phase by that time (Fig. 2A). In both Fig. 2B and 2C, histone H1 does not approach full phosphorylation in calyculin A-induced PCC until T=16 or 20 hrs, even though 80% of the cells are already in S-phase by T=12 hrs (Fig. 2A).

The lack of H1 phosphorylation in early S-phase PCC (Fig. 2B and C) may appear to be inconsistent with the results in Fig. 1E. Fig. 1E shows that H1 is almost fully phosphorylated when PCC is induced in thymidine-treated cells, many of which should be in early S-phase (Xeros, 1962; Merrill, 1998). This apparent discrepancy is explained by the fact that chemical treatments only produce cell cycle synchrony with respect to some processes, not all (Bergeron, 1971; Cooper, 2003). Although thymidine may block DNA synthesis in early S-phase, other S-phase processes, such as cyclin synthesis, likely continue (Heintz et al., 1983).

The results in Fig. 2 suggest that histone H1 is not phosphorylated in calyculin A-induced PCC until the cells have progressed at least four hours into S-phase. They also confirm that butyrate arrests HeLa cells in G_1_-phase and that butyrate-treated cells can undergo PCC. We conclude that any PCC induced by calyculin A-treatment in G_1_ and early S-phase will lack histone H1 phosphorylation.

### Calyculin A induces PCC in early G_1_-phase cells

For the experiments shown in Fig. 1F and Fig. 2, G_1_-phase cells were obtained by treating cultures with sodium butyrate. To further investigate G_1_-PCC, early G_1_-phase cells were produced by inducing exit from mitotic arrest.

Metaphase-arrested HeLa cells can be triggered to exit mitosis and enter G_1_ (though without chromosome segregation or cytokinesis) by inactivating Cdk1/cyclin B or by inhibiting it using cell-permeable Cdk1 inhibitors (Paulson, 2007, 2024; Keaton et al., 2023). For example, treatment of metaphase-arrested HeLa cells with staurosporine, an inhibitor of Cdk1 and other protein kinases (Gadbois et al., 1992), causes the cells to decondense their chromosomes, reassemble nuclear envelopes, and dephosphorylate histone H1 without undergoing cytokinesis (Th’ng et al., 1994; Hall et al., 1996; Paulson et al., 1996). Here, we used the Cdk1 inhibitor roscovitine (Meijer et al., 1997).

The experiment shown in Fig. 3 used a metaphase-arrested HeLa culture (mitotic index, 96%). One portion of the culture was not treated with roscovitine and remained in mitosis for the duration of the experiment (○---○). The rest of the cells were treated with 200 µM roscovitine for 50 min (●··─··●). By T=50 min the treated cells had decondensed their chromosomes and reassembled nuclei.

**Fig. 3.**
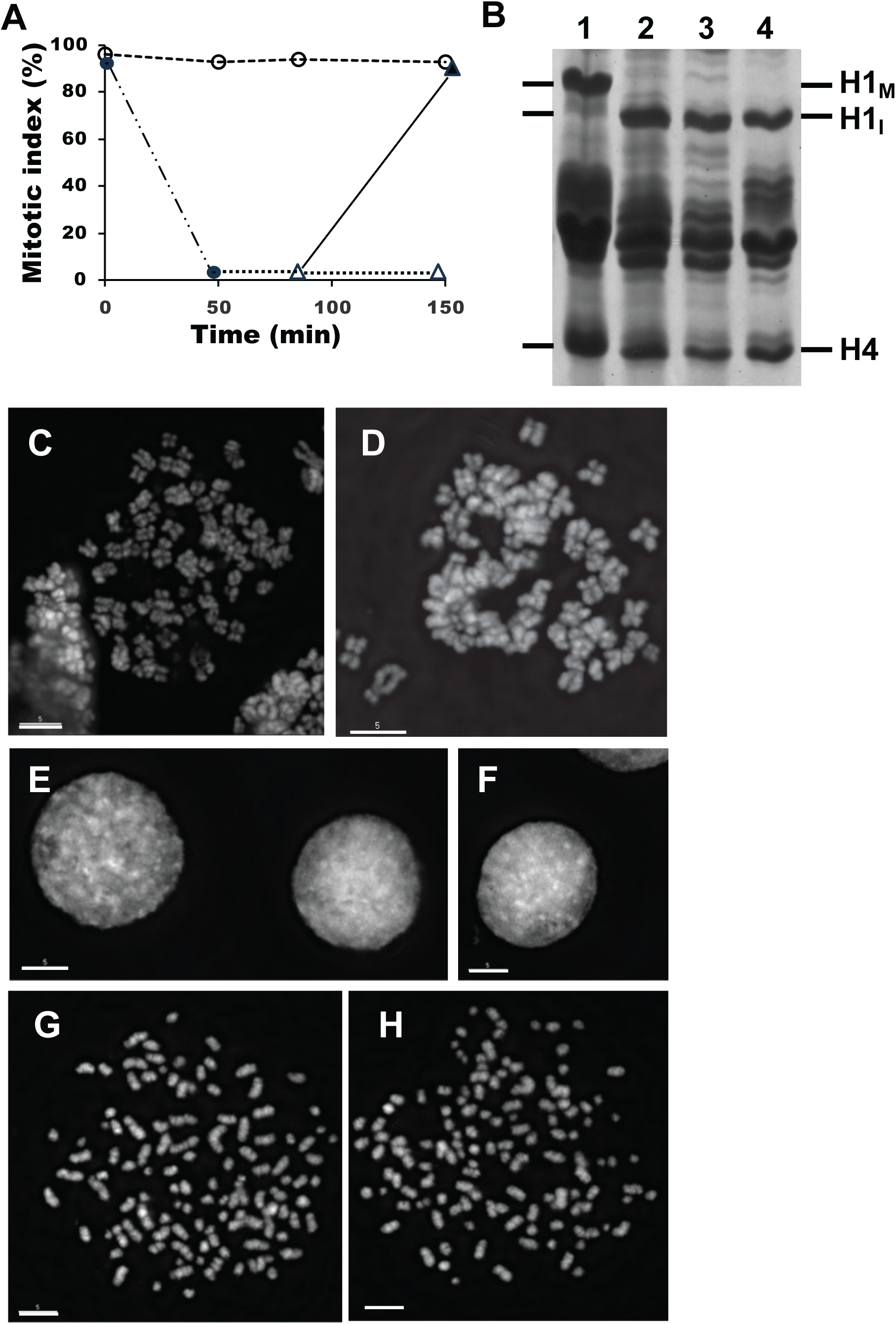
Roscovitine treatment of metaphase-arrested HeLa cells induces exit from mitosis and subsequent calyculin A treatment induces G_1_-PCC with single chromatids and no histone H1 phosphorylation. (A) The “mitotic index” (percentage of cells with condensed chromosomes and no nuclear envelopes) as a function of time after various treatments. Aliquots of a metaphase-arrested culture (mitotic index, 96%) were either incubated with no treatment (○---○) or treated with 200 µM roscovitine for 50 min (●··─··●). At T=50 min, roscovitine was removed from the treated cells, some of which were incubated without further treatment until T=150 min (Δ···Δ) while others were treated with 100 nM calyculin A at T=85 min to induce PCC (▴—▴); (B) Histones extracted from samples in (A) and analyzed by acid-urea gel electrophoresis: Metaphase-arrested cells at T=0 min (lane 1); cells at T=50 min after roscovitine treatment (lane 2); cells after roscovitine removal and further incubation until T=150 min (lane 3); and cells after roscovitine removal, treatment with calyculin A at T=85 min, and further incubation until T=150 min (lane 4). (C-H) Examples of fixed cells examined by fluorescence microscopy with DAPI stain. (C, D) Metaphase-arrested cells at T=0 min corresponding to Panel B, lane 1. The condensed chromosomes consist of paired chromatids. (E, F) Cells at T=50 min after treatment with 200 µM roscovitine, corresponding to Panel B, lane 2. Chromosomes have decondensed and nuclear envelopes have reassembled. (G, H) Roscovitine-treated cells (as in E and F) that were pelleted, washed, resuspended in fresh medium and treated with calyculin A, corresponding to Panel B, lane 4. The condensed chromosomes (G_1_-PCCs) consist of single chromatids and lack histone H1 phosphorylation. Scale bars represent 5 µm.

After T=50 min, the roscovitine-treated cells were pelleted, washed, and resuspended in fresh medium. One portion was incubated without further treatment and those cells remained in interphase until T=150 min (Δ···Δ). Another portion was treated with 100 nM calyculin A from T=85 to 150 min (▴—▴). G_1_-PCC was observed in nearly all of those cells (Fig. 3A).

Fig. 3B shows that histone H1 was phosphorylated in the metaphase-arrested cells (Fig. 3B, lane 1) but dephosphorylated during the roscovitine treatment (lane 2). When the roscovitine was removed and those cells were incubated until T=150 min without further treatment, H1 remained unphosphorylated (lane 3). However, even after treatment with calyculin A to efficiently induce G_1_-PCC, H1 was not phosphorylated (Fig. 3B, lane 4).

Fluorescence microscopy of fixed, spread DAPI-stained metaphase-arrested cells at the start of the experiment shows condensed duplex chromosomes (Fig. 3C and D).

After roscovitine treatment for 50 min, only nuclei are seen (Fig. 3E and F), indicating exit from metaphase-arrest to G_1_-phase. After further treatment with 100 nM calyculin A for 65 min, nuclear envelopes break down again, chromosomes condense, and G_1_-PCC are seen (Fig. 3G and H).

These results demonstrate that calyculin A-treatment of G_1_-phase cells induces condensed chromosomes (G_1_-PCC) without histone H1 phosphorylation. Similar results have been obtained with HeLa cells that exited metaphase-arrest due to removal of nocodazole, and with mouse FT210 cells, which have temperature-sensitive Cdk1 (Mineo et al., 1986; Th’ng et al., 1990), that were induced to leave metaphase-arrest by heat treatment (Paulson and Vander Mause, 2013). In all cases, treatment with calyculin A brings about nuclear envelope breakdown and chromosome condensation (G_1_-PCC), but histone H1 is not phosphorylated.

Interestingly, while metaphase cells contain paired chromatids (Fig. 3C and D), chromosomes in the cells with G_1_-PCC are characterized by single chromatids, and there are approximately twice as many of them as duplex chromosomes in metaphase (Fig. 3G and H). This shows that triggering mitotic exit in prometaphase-arrested cells by artificial inactivation of Cdk1/cyclin B leads to sister chromatid separation. Since mitotic exit was induced without cytokinesis, these G_1_ cells are now tetraploid. A similar phenomenon is observed when FT210 cells (with temperature-sensitive Cdk1) are induced to exit metaphase-arrest by heat treatment (Paulson, 2007).

### Establishing cell lines for FLIM-FRET

Extensive phosphorylation of histone H1 coincides with mitosis (e.g., Gurley et al., 1978) and was traditionally proposed to play a role in mitotic chromosome condensation (Bradbury et al., 1973; Th’ng et al., 1994). However, the results presented above suggest that chromosome condensation is independent of H1 phosphorylation.

Still, it is possible that while phosphorylation of H1 is not required for the overall morphology of condensed chromatids, it may make the chromatin more compact. We therefore asked whether chromatin is compacted to the same extent in mitotic chromosomes and in G_1_-PCC.

This question was addressed using FLIM-FRET technology, which can quantitate the extent of chromatin compaction by measuring energy transfer between fluorescent proteins fused to histone H2B (Llères et al., 2009). The efficiency of energy transfer indicates how close the donor and acceptor proteins are to one another.

Green and red fluorescent proteins were fused to the C- and N- termini of H2B, respectively, allowing them to interact between neighboring nucleosomes, but not within the same nucleosome (Llères et al., 2009). Experimental and control constructs were cloned into the piggyback transposase vector to boost the expression of ectopically expressed histone H2B, which is difficult to overexpress relative to the endogenous control (Llères et al., 2009; Dupont et al., 2023). Cell lines were established in a HeLa CDK1^as^ background (Rata et al., 2018) to allow efficient synchronization using 1NM- PP1.

Expressed H2B-mNeonGreen localized to the nucleus and mitotic chromosomes (Fig. 4A, 1, 2, 3) as did mScarlet-I-H2B (Fig. 4A, 1). Ectopically expressed freely floating mScarlet-I in a control cell line localized to the cytoplasm of mitotic cells and exhibited both nuclear and cytoplasmic localization in interphase cells (Fig. 4A, 2). In a positive control cell line expressing mNeonGreen-linker-mScarlet-I (which produces constitutively active FRET), the fusion protein localized to both nuclei and cytoplasm in interphase and was not enriched on chromosomes in mitosis (Fig. 4A, 4).

**Fig. 4.**
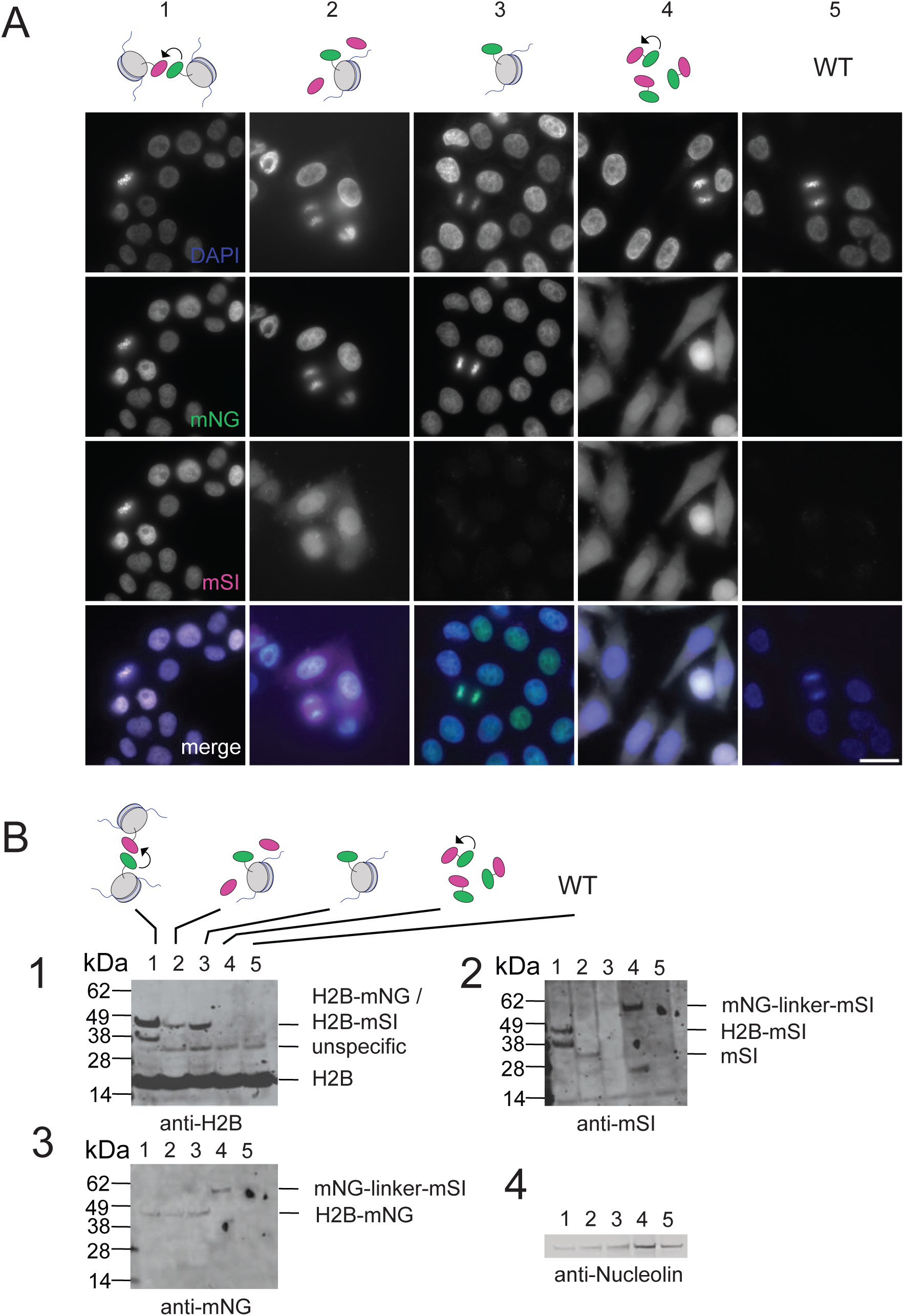
Characterization of cell lines used for FLIM-FRET. (A) Fluorescence images of the cell lines used for FLIM-FRET, expressing (1) H2B-mNeonGreen and mScarlet-I- H2B; (2) H2B-mNeonGreen and free mScarlet-I; (3) H2B-mNeonGreen; (4) mNeonGreen-linker-mScarlet-I; and (5) untransfected (wild-type). Fluorescence images show DAPI staining, mNeonGreen (mNG), mScarlet-I (mSI), and all three (merge). The exposure of the images was adjusted individually. For channels of cell lines not expressing certain fluorescent proteins, the scale was adjusted at maximum channel exposure, represented in the panel. (B) Expression of exogenous proteins in the five cell lines analyzed by western blots using primary antibodies against H2B (1), mScarlet-I (2), mNeonGreen (3) and nucleolin (loading control) (4). Labeling as in (A). Brightness and contrast were adjusted for individual panels.

The expression of exogenous proteins in these cell lines was confirmed by western blotting (Fig. 4B). All overexpressed proteins were detected at the expected size, and H2B-mNeonGreen fusions showed comparable expression between the different cell lines (Fig. 4B3).

### G_1_-PCC induction by calyculin A in adherent HeLa CDK1^as^ cells

G_1_-PCC were induced in HeLa CDK1^as^ cells (Rata et al., 2018) by treatment of G_1_ cells with calyculin A. The synchronization procedure to obtain G_1_ cells (Fig. 5A) included a 2.5 mM thymidine block for 18 hrs. After thymidine wash-out, low amounts of thymidine (TdR) and deoxycytidine (CdR) were added to the medium to help cells recover from depletion of pyrimidines (Heintz et al., 1983), and the cells were treated with 1NM-PP1 for 8 hrs to block them in G_2_ phase. An hour after the cells were released into mitosis by removal of 1NM-PP1, mitotic cells were shaken off and allowed to re-adhere, with thymidine present in the medium to prevent entry into S-phase. After 3 hrs, calyculin A was added for 2 hrs (Fig. 5A) to induce PCC. In controls, asynchronous cultures were blocked with nocodazole for 14 hrs.

**Fig. 5.**
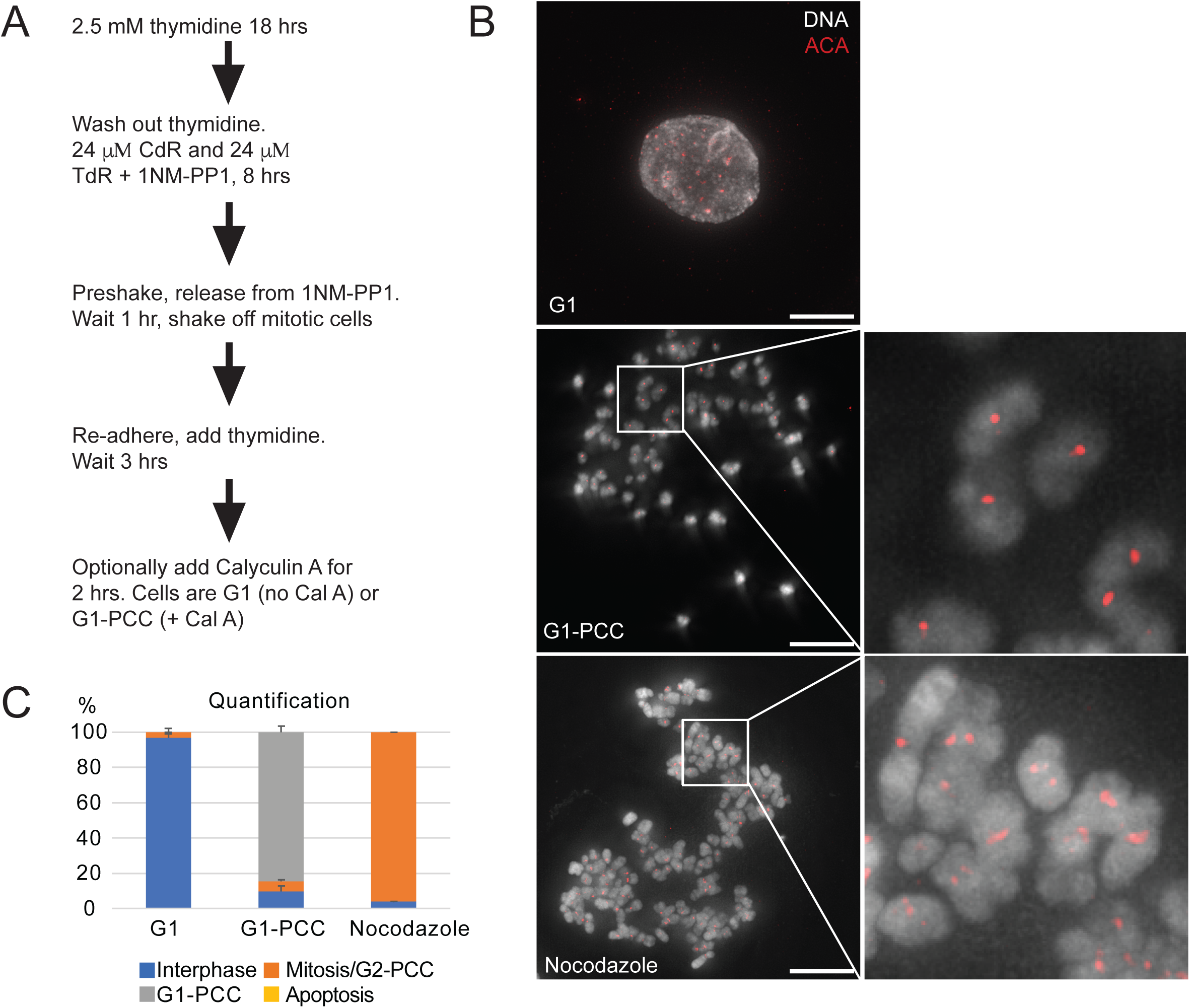
Cell synchronization in G_1_, G_1_-PCC and mitosis for the FLIM experiment. (A) The scheme of cell synchronization before G_1_-PCC induction. (B) Methanol/acetic acid fixation and immunofluorescence with anti-centromere antibodies (ACA) of G_1_(upper), G G_1_1-PCC (middle) and nocodazole-treated cells (lower). Enlarged insets of G_1_-PCC and metaphase chromosomes are shown on the right. Images of G_1_-PCC spreads and G_1_ cells were taken in one experiment (3 replicates), metaphase spreads of cells arrested in nocodazole were imaged in a separate experiment (2 replicates). Brightness and contrast of the centromeres on condensed chromosomes were adjusted to comparable levels. Maximum intensity projections of DAPI and ACA channels are shown. Scale bars represent 10 µM. (C) Quantification of phenotypes based on the presence of nuclei versus chromosomes and on the number of ACA dots on the chromosomes. A minimum of 200 cells or spreads were counted in each condition. G_1_-PCC spreads and G_1_cells were counted in one experiment (3 replicates), metaphase spreads of cells arrested in nocodazole were counted in a separate experiment (2 replicates).

Methanol-acetic acid fixation and immunofluorescence with anti-centromere antibodies (ACA) (Earnshaw and Rothfield, 1985) showed single centromeres on G_1_- PCC chromosomes, compared to two centromeres on mitotic chromosomes (one centromere per chromatid). (Fig. 5B). Quantification of the chromosome morphology confirmed that we had obtained synchronous populations of G_1_ cells, G_1_-PCC, and mitotic cells (Fig. 5C).

### No difference in compaction between G_1_-PCC and mitotic chromosomes

The established cell lines were synchronized as described above and G_1_, G_1_-PCC, and mitotic cells were imaged for FLIM-FRET measurements in 3 biological replicates. When FRET occurs, energy transfer from the donor to the acceptor reduces the fluorescence lifetime of the donor. As expected, the mNeonGreen lifetime in the positive control cell line expressing mNeonGreen-linker-mScarlet-I was reduced compared to the cell line expressing donor only (H2B-mNeonGreen) (Fig. 6A1; Table S2). In the cell line expressing both H2B-donor and acceptor-H2B fusions (H2B-mNeonGreen and mScarlet-I-H2B), the lifetime of the donor was reduced compared to donor only (H2B- mNeonGreen) (Fig. 6A2; Fig. S1A, B), or donor only with freely floating mScarlet-I (Fig. 6A2; Table S2). Background lifetimes with the non-fluorescent parental cell line (“wild type”, WT) were randomly distributed and close to 0 (Fig. S1C).

**Fig. 6.**
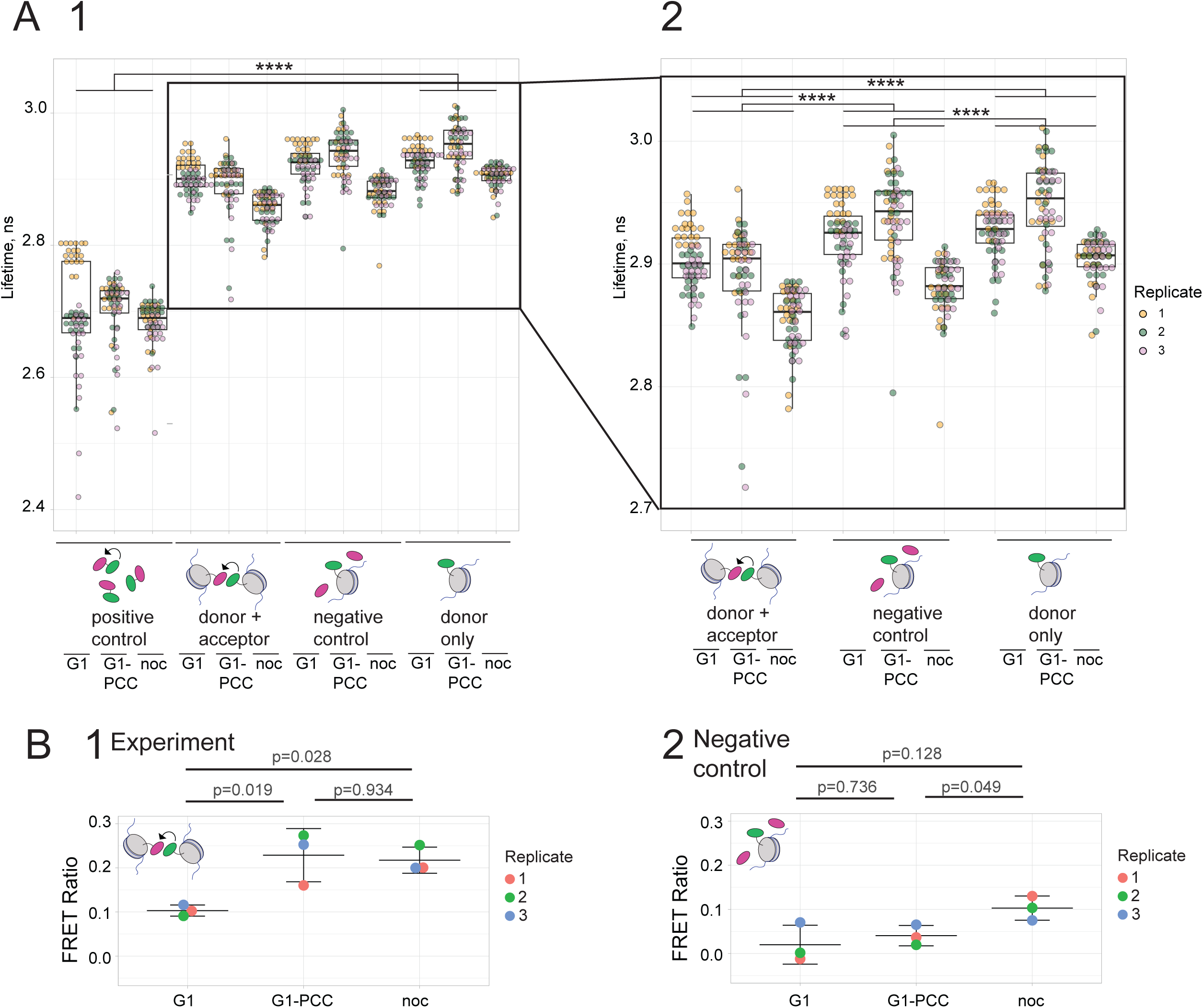
G_1_-PCC and mitotic chromosomes display no differences in chromatin compaction as detected by FLIM-FRET. (A1) Distributions of lifetimes in the FLIM-FRET experiment. From left to right, cells expressing: mNeonGreen-linker-mScarlet-I (positive control); H2B-mNeonGreen and mScarlet-I-H2B (donor + acceptor); H2B- mNeonGreen and free mScarlet-I (negative control); and H2B-mNeonGreen (donor only). For each cell line, results for G_1_, G_1_-PCC, and nocodazole-arrested cells are shown from left to right. (A2) Magnified view of (A1) for all cell lines except the positive control. In A1 and A2, separate conditions and replicates were imaged on differing days. For statistical comparison, linear mixed effect models were fit for log(Lifetime). Cell line and condition were fit as fixed effects and slide (single replicate of a particular condition, i.e. the day of experiment) as a random effect. In the zoomed in plot (A2), the p-value adjustment was performed with the Benjamini Hochberg procedure. (B) Ratios of FRET efficiencies. FRET efficiencies were calculated as (1 – τ_DA_/τ_D_) using average lifetimes τ_DA_ for mNeonGreen-linker-mScarlet-I (positive control), H2B-mNeonGreen and mScarlet-I-H2B (donor + acceptor), and H2B-mNeonGreen and free mScarlet-I (negative control), and the average lifetime τ_D_ for H2B-mNeonGreen (donor only). FRET ratios were then calculated for (B1) H2B-mNeonGreen and mScarlet-I-H2B (donor + acceptor), and (B2) H2B-mNeonGreen and free mScarlet-I (negative control). This was done by dividing the FRET efficiencies of those cells by the FRET efficiency of the positive control, always using values from replicates processed on the same occasion under the same conditions. In each case, the data was analyzed with one-way ANOVA followed by the post-hoc Tukey test. The normality of distributions of residuals and homogeneity of variances were confirmed.

The mean lifetime of the positive control was variable between replicates (Fig. 6A1) performed on different days, presumably due to differences in adjustments to the microscope, laser, and detector. However, on a given occasion, replicates for all five cell lines were processed within a short time window, 45 min apart, using the same equipment and under the same conditions. We therefore compared compaction in G_1_, G_1_-PCC and nocodazole-arrest by computing the ratios of the FRET efficiencies to the FRET efficiency of the positive control for each replicate. These “FRET ratios” served to normalize the FRET efficiencies.

For each replicate, the FRET efficiency for donor plus acceptor (H2B- mNeonGreen/mScarlet-I-H2B) was divided by the FRET efficiency for the positive control (mNeonGreen-linker-mScarlet-I) of that replicate. These ratios are given in Fig.6B1 and Table 1. We also divided the FRET efficiency for the negative control (H2B-mNeonGreen/mScarlet-I) by the FRET efficiency for the positive control (mNeonGreen-linker-mScarlet-I) and these ratios are also given in Fig.6B2 and Table 1. The donor plus acceptor vs positive control FRET ratios were similar between replicates for G_1_ cells, but the ratios were somewhat variable for condensed chromosomes (Fig. 6B1). This might suggest a stochastic component in the level of condensed chromosome compaction.

**Table 1:**
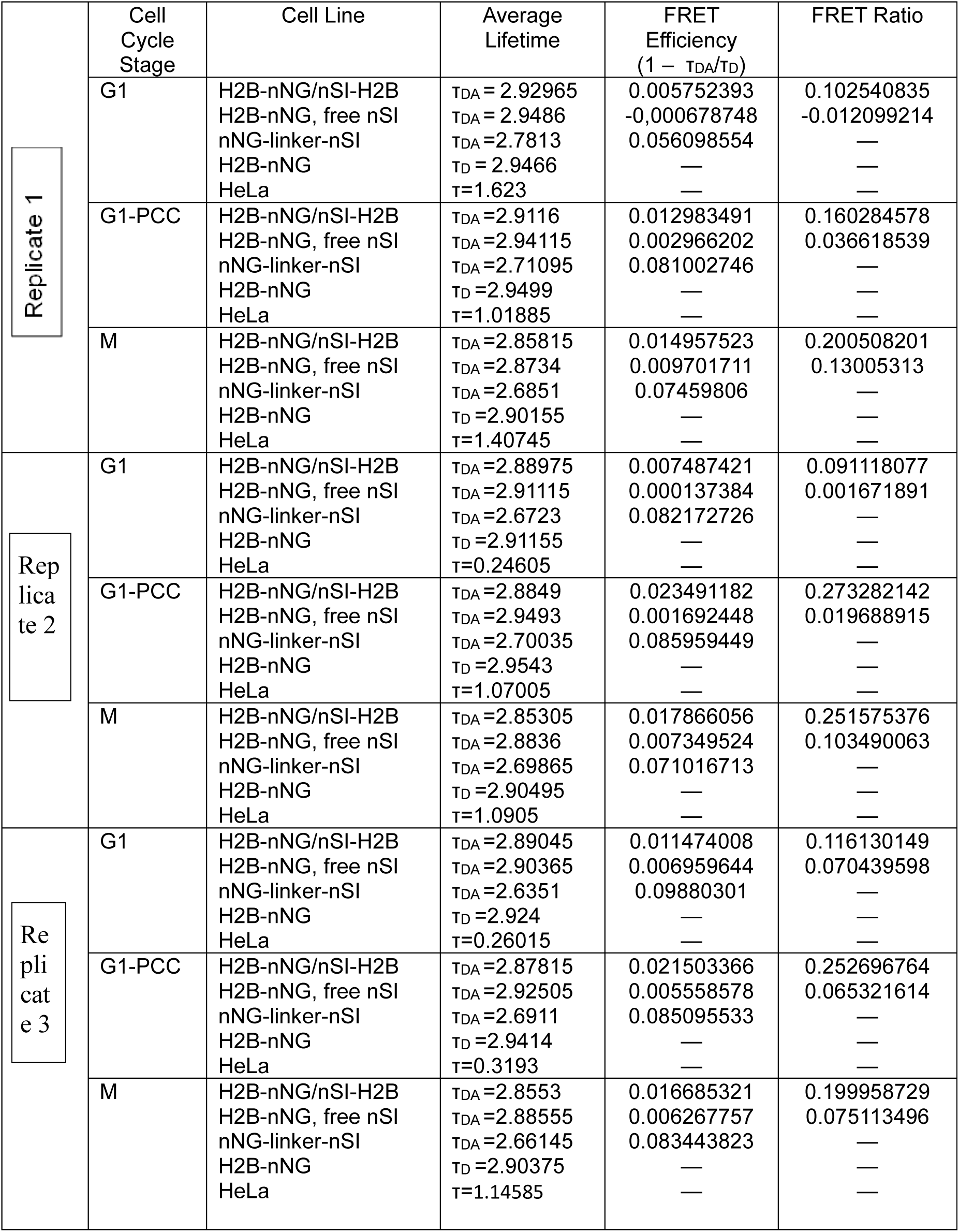
Average lifetimes, FRET efficiencies and FRET efficiency ratios for each experiment.

Most importantly, however, we observed no significant difference in compaction between G_1_-PCC and mitotic chromosomes (p=0.934), and both were more compact than G_1_ chromatin (p=0.019 and p=0.028) (Fig. 6B1). On the other hand, for the negative control vs positive control FRET ratios (Fig. 6B2), no differences between G_1_, G_1_-PCC and nocodazole arrest were detected.

## Discussion

### Calyculin A induces premature chromosome condensation in G_1_- and early S-phase HeLa cells without histone H1 phosphorylation

Our results confirm previous observations that premature chromosome condensation (PCC) can be induced in vertebrate cells at any point in interphase by treatment with calyculin A, an inhibitor of protein phosphatases PP1 and PP2A (e.g., Gotoh et al., 1995; Coco-Martin and Begg, 1997; Alsbeih and Raaphorst, 1999; Kanda et al., 1999; Gotoh, 2009). However, we find that PCC induced in G_1_- or early S-phase cells is not accompanied by histone H1 phosphorylation. Despite this, chromosomes in G_1_-PCC closely resemble metaphase chromatids, providing the first clear demonstration of mitotic-like chromosome formation without H1 phosphorylation.

Although we have not determined cyclin levels in this study, it appears that histone H1 phosphorylation in PCC correlates with the presence of cyclin B. In normal mitosis, H1 is phosphorylated by Cdk1/cyclin B (e.g., Labbé et al., 1989; Langan et al., 1989); indeed, H1 was one of the canonical substrates used in the isolation of the critical cell cycle kinase MPF (Maturation Promoting Factor), which turned out to be Cdk1/Cyclin B (Lohka et al., 1988; Gautier et al., 1988; Gautier et al., 1990). However, cyclin B is proteolytically degraded during mitotic exit and is not resynthesized until the next S-phase (Frisa and Jacobberger, 2009; Huang et al., 2001; Fung and Poon, 2005; Lindqvist et al., 2009). Thus, the timing of cyclin B availability fits with our observations that H1 is phosphorylated in G_2_- and late S-phase PCC, apparently to the same extent as in mitosis, but not in G_1_- or early S-phase PCC.

### Chromatin in G_1_-PCC is as compact as in metaphase chromosomes, despite lack of histone H1 phosphorylation

Contrary to earlier proposals that H1 phosphorylation is responsible for chromosome condensation (Bradbury et al., 1973; Th’ng et al., 1994), our results clearly demonstrate that H1 phosphorylation is not essential for the formation of mitotic-like chromosomes. Although chromosomes in early G_1_-phase PCC completely lack phosphorylated H1, in the light microscope they look just like metaphase chromosomes, except that they consist of single chromatids.

However, ordinary light microscopy cannot show whether H1 phosphorylation affects the tightness of chromatin packing within condensed chromosomes. To measure chromatin compaction, we used FLIM-FRET (Llères et al., 2009) to compare the separation between donor and acceptor H2B molecules in G_1_ cells, G_1_-PCC, and nocodazole-arrested cells. Our results show no difference in chromatin compaction between G_1_-PCC and mitotic chromosomes, but both are more compact than chromatin in G_1_-phase nuclei.

### Which protein kinases are involved in G_1_-PCC?

The occurrence of chromosome condensation and chromatin compaction in G_1_- PCC without histone H1 phosphorylation raises interesting questions concerning the protein kinases involved in G_1_-PCC, the process of condensed chromosome formation, and the mechanism of chromatin compaction.

Cdk1^as^ cells are efficiently arrested in late G_2_ by specific inhibition of Cdk1 with 1NM-PP1 (Hochegger et al., 2007; Rata et al., 2018; Samejima et al., 2018; Samejima et al., 2022; Gibcus et al., 2018), showing that normal mitotic entry absolutely requires Cdk1/Cyclin B. Cdk1 is required for nuclear envelope breakdown at mitosis (e.g., de Castro et al., 2017) and for activation of various condensin subunits (Kimura et al., 1998, 1999; Shintomi et al., 2015; Holt et al., 2009; Tane et al., 2022; Yoshida et al., 2024).

Calyculin A-induced G_1_-PCC presumably involves protein phosphorylation, since calyculin A is a protein phosphatase inhibitor, but it does not involve Cdk1/cyclin B. The present experiments and their predecessors (e.g., Gotoh et al., 1995; Coco-Martin and Begg, 1997; Alsbeih and Raaphorst, 1999; Kanda et al., 1999; Gotoh, 2009) show that mitotic-like nuclear envelope breakdown and chromosome formation occur in G_1_-PCC, though they normally involve Cdk1, and the absence of Cdk1/cyclin B involvement in G_1_-PCC is confirmed by the lack of phosphorylated H1. Cdk1 must be inactive in G_1_- phase to allow for CENP-A deposition, which occurs in G_1_ but is blocked by Cdk1 phosphorylation of MIS18BP1 (Jansen et al., 2007; Silva et al., 2012; Spiller et al., 2017; Pan et al., 2017).

Whatever protein kinases are involved in induction of G_1_-PCC, their effects follow from calyculin A inhibition of protein phosphatases. Either the relevant kinases are held in an inactive state by a protein phosphatase, or they are constitutively active but their effects are countered by phosphatase activity. Inhibition of PP1 in G_1_ could trigger a non-canonical, Cdk1-independent pathway of mitotic entry.

The protein kinases at work in calyculin A-induced G_1_-PCC could include Plk1, which promotes chromosome condensation by condensin (Abe et al., 2011) and regulates Mis18BP1 activity during G1 (Parashara et al., 2024; Conti et al., 2024). Another candidate is Aurora B. Phosphorylation of histone H3 on serine 10 is primarily regulated by Aurora B and PP1 and has been correlated with mitotic chromosome condensation (Gurley et al., 1978; Hendzel et al., 1997; Bradbury, 1992; Kaszas and Cande, 2000; Wei et al., 1999; Ajiro et al., 1996; Hsu et al., 2000; Murnion et al., 2001). Aurora B is also involved in condensin I loading onto chromosomes (Lipp et al., 2007). Alternatively, G_1_- PCC could involve Cdk2 (though that may require cyclin B; Lau et al., 2021), Cdk4/6, or other protein kinases.

This will be interesting to explore in future studies because the protein kinases involved in G_1_-PCC may also have roles in normal mitosis. Comparing the phosphoproteins in G_1_-PCC and mitotic cells may provide clues. Another approach will be to block the action of various kinases during the induction of PCC and study the effects on chromosome condensation, chromatin compaction, and nuclear envelope breakdown. This could be done using gene knockdowns, or kinase inhibitors such as ZM447439 which inhibits Aurora B (Gadea and Ruderman, 2005). Such experiments may be more convenient using G_2_- rather than G_1_-PCC. If Cdk1^as^ cells arrested in G_2_ with 1NM-PP1 are treated with calyculin A, they also efficiently undergo PCC (Ustun et al., in preparation).

### What drives mitotic-like chromosome formation (PCC) during G_1_ phase?

It is not surprising that the overall morphology of mitotic chromosomes does not require H1 phosphorylation, since mitotic-like chromosome structures can assemble *in vitro* without any histone H1 and with few core histones (Shintomi et al., 2015; Shintomi et al., 2017). On the other hand, a more recent study suggests that H1 might interfere with chromatin binding by condensins and DNA topoisomerase II to help determine the morphology of mitotic chromosomes (Choppakatla et al., 2021).

Mitotic chromosome formation is primarily driven by the SMC (Structural Maintenance of Chromosomes) proteins condensin II and condensin I (Hirano, 2016; Yatskevich et al., 2019; Hoencamp and Rowland, 2023; Naumova et al., 2013; Shintomi and Hirano, 2011; Ono et al., 2003; Paulson et al., 2021). Condensin II is normally found in the nucleus where Mcph1 (Trimborn et al., 2006; Yamashita et al., 2011; Houlard et al., 2021) prevents it from acting on chromosomes until Mis18BP1 displaces the Mcph1 (Borsellini et al., 2025). A broad consensus suggests that condensin II activity involves the extrusion of chromatin loops to a size of about 400 kb (Nasmyth, 2001; Alipour and Marko, 2012; Goloborodko et al., 2016; Dekker and Mirny, 2024; Samejima et al., 2025; Ganji et al., 2018; Pradhan et al., 2022; Davidson et al., 2019; Golfier et al., 2020; Barth et al., 2025; Lee et al., 2025), while Condensin I forms nested loops of about 80 kb, further condensing the structure (Gibcus et al., 2018). Although some workers propose that loop extrusion is not essential (Gerguri et al., 2021; Forte et al., 2024; Forte et al., 2025; Uhlmann, 2025), the bulk of evidence supports the loop extrusion theory.

Condensin I and II action results in a bottlebrush-like array of loops with the characteristic rod-like shape of a chromatid, within which condensin II appears to follow a disorderly helical path (Gibcus et al., 2018; Samejima et al., 2025; Chu et al., 2020).

A previous study argues (Zhang et al., 2016), that the formation of morphologically normal chromatids in G_1_-PCC is also driven by condensins. But how condensins are regulated in this case is a question that requires further study. As previously mentioned, numerous studies report activation of various condensin subunits by Cdk1 (Kimura et al., 1998; Kimura et al., 1999; Shintomi et al., 2015; Holt et al., 2009; Tane et al., 2022; Yoshida et al., 2024), but which kinase(s) might be involved will require further study.

### What causes chromatin compaction in G1-PCC and mitosis?

The cause of chromatin compaction in mitotic chromosomes is not known, but we have reason to believe that the same cause operates in G_1_-PCC. Several possibilities can therefore be tested using calyculin A-induced PCC and FLIM-FRET. For example, condensins I and II may contribute to chromatin compaction via DNA supercoiling during chromatin loop extrusion (e.g., St. Pierre et al., 2009; Janissen et al., 2024). This could be tested by inducing PCC in normal cells and in cells where SMC2 has been eliminated and using FLIM-FRET to compare chromatin compaction. Without SMC2, and therefore without condensins I and II, morphologically normal chromosomes do not form, but the chromatin is still highly condensed as seen by light microscopy (Hudson et al., 2003; Samejima et al., 2018).

Histone deacetylation may also promote mitotic chromatin compaction (e.g., Wilkins et al., 2014). To test this, cells could be treated with histone deacetylase inhibitors in advance of mitosis (e.g., Patzlaff et al., 2010) or in advance of calyculin A treatment. FLIM-FRET could then be used to compare chromatin compaction in cells with and without histone hyperacetylation.

Other post-translational modifications of histones may also contribute to mitotic chromatin compaction (Andonegui-Elguera et al., 2022). For example, Zhiteneva et al. (2017) found that chromatin arrays assembled from mitotic core histones lacking H1 were more prone to aggregation than identical arrays assembled from interphase histones. Of course, to elucidate the roles of various modifications, it will be necessary to find ways to artificially control them. One must also bear in mind that events such as chromatin compaction or nuclear envelope breakdown could result from the additive effects of multiple factors, complicating these types of experiments.

## Conclusion

In summary, our results indicate that mitotic chromosome morphology and chromatin compaction do not require histone H1 phosphorylation. The function of mitotic H1 phosphorylation remains unknown, but it may play a role in H1 mobility (Contreras et al., 2003), the kinetics of packaging, or the stability of condensed chromosomes, and possibly it could affect nucleosome arrangement in ways that our FLIM-FRET experiments cannot detect.

Our results also suggest that calyculin A-induced G_1_-PCC could be useful in experiments designed to identify factors involved in condensed chromosome formation, chromatin compaction, nuclear envelope breakdown, and other mitotic events.

Information obtained in this way may provide clues to the enzymes and mechanisms involved in normal mitosis.

## Acknowledgements

We thank Masato T. Kanemaki for sharing the piggyback plasmid with blastocidin resistance, as well as the laboratories of Alistair McCormick, Julie Welburn, and Patrick Heun at the University of Edinburgh for gifting plasmids containing fluorescent proteins and histone H2B. We also thank Lorenza Di Pompeo for advice on establishing cell lines, and Itaru Samejima for assistance with HeLa S3. We acknowledge Antoni Sieminski and the Centre for Statistics Drop-in Clinic at the University of Edinburgh for help with statistical analysis of mNeonGreen lifetimes. This work was supported by grant S352 (to J.R. Paulson) from the University of Wisconsin-Oshkosh Faculty Development Board, and also by grants R15 GM39915 and R15 GM46040 (to J. R. Paulson) and grant R01 CA69008 (to W. C. Earnshaw and S. H. Kaufmann) from the National Institutes of Health, U.S. Public Health Service. B. Prevo was supported by a Sir Henry Wellcome Postdoctoral Fellowship (no. 215925) sponsored by W. C. Earnshaw. The W.C. Earnshaw lab was also supported by the Wellcome Principal Research Fellowship no. 107022.

## Supplementary Material

Table S1

Table S2

Fig. S1

## References

1. Abad MA, Ruppert JG, Buzuk L, Wear M, Zou J, Webb KM, Kelly DA, Voigt P, Rappsilber J, Earnshaw WC, Jeyaprakash AA (2019) Borealin–nucleosome interaction secures chromosome association of the chromosomal passenger complex. J Cell Biol 218:3912–3925. PMID: 31570499. doi: 10.1083/jcb.201905040

2. Abe S, Nagasaka K, Hirayama Y, Kozuka-Hata H, Oyama M, Aoyagi Y, Obuse C, Hirota T (2011) The initial phase of chromosome condensation requires Cdk1- mediated phosphorylation of the CAP-D3 subunit of condensin II. Genes Dev 25:863–74. PMID: 21498573. doi: 10.1101/gad.2016411.

3. Ajiro K, Yoda K, Utsumi K, Nishikawa Y (1996) Alteration of cell cycle-dependent histone phosphorylations by okadaic acid. Induction of mitosis-specific H3 phosphorylation and chromatin condensation in mammalian interphase cells. J Biol Chem 271:13197–201. PMID: 8662672. doi: 10.1074/jbc.271.22.13197.

4. Akera T, Watanabe Y (2016) The spindle assembly checkpoint promotes chromosome bi- orientation: A novel Mad1 role in chromosome alignment. Cell Cycle 15(4):493–497. PMID: 26752263. doi: 10.1080/15384101.2015.1128596

5. Alipour E, Marko JF (2012) Self-organization of domain structures by DNA-loop-extruding enzymes. Nucleic Acids Res 40:11202–12. PMID: 23074191. doi: 10.1093/nar/gks925.

6. Alsbeih G, Raaphorst GP (1999) Differential induction of premature chromosome condensation by calyculin A in human fibroblast and tumor cell lines. Anticancer Res. 19: 903–908. PMID: 10368632.

7. Alvarez LAJ, Widzgowski B, Ossato G, van den Broek B, Jalink K, Kuschel L, Roberti MJ, Hecht F (2019) Application Note: SP8 FALCON: a novel concept in fluorescence lifetime imaging enabling video-rate confocal FLIM. Nature Methods. https://www.nature.com/articles/d42473-019-00261-x (Accessed 2024-11-12)

8. Andonegui-Elguera MA, Cáceres-Gutiérrez RE, López-Saavedra A, Cisneros-Soberanis F, Justo-Garrido M, Díaz-Chávez J, Herrera LA (2022) The roles of histone post-translational modifications in the formation and function of a mitotic chromosome. Int J Mol Sci 23: 8704. PMID: 27818790. doi: 10.3390/ijms23158704

9. Barth R, Davidson IF, van der Torre J, Taschner M, Gruber S, Peters J-M, Dekker C (2025) SMC motor proteins extrude DNA asymmetrically and can switch directions. Cell 188:749–763.e21. PMID: 39824185. doi: 10.1016/j.cell.2024.12.020.

10. Bergeron JJM (1971) Different effects of thymidine and 5-fluorouracil 2′-deoxyriboside on biosynthetic events in cultured P815Y mast cells. Biochem J 123: 385–390. PMID: 4256660. doi: 10.1042/bj1230385

11. Bezrookove V, Smits R, Moeslein G, Fodde R, Tanke HJ, Raap AK, Darroudi F (2003) Premature chromosome condensation revisited: a novel chemical approach permits efficient cytogenetic analysis of cancers. Genes Chromosomes Cancer 38: 177–186

12. Bishop AC, Ubersax JA, Petsch DT, Matheos DP, Gray NS, Blethrow J, Shimizu E, Tsien JZ, Schultz PG, Rose MD, Wood JL, Morgan DO, Shokat KM (2000) A chemical switch for inhibitor-sensitive alleles of any protein kinase. Nature 407: 395–401.

13. Borsellini A, Conti D, Cutts E, Harris RJ, Walstein K, Graziadei A, Cecatiello V, Aarts TF, Xie R, Ruzi AM, Sen S, Hoencamp C, Pleuger R, Ghetti S, Oberste-Lehn L, Pan D, Bange T, Haarhuis JHI, Perrakis A, Brummelkamp TR, Rowland BD, Musacchio A, Vannini A (2025) Condensin II activation by M18BP1. bioRxiv. 2024.05.02.592151. 10.1101/2024.05.02.592151.

14. Bradbury EM (1992) Reversible histone modifications and the chromosome cell cycle. Bioessays 14:9–16. PMID: 1312335. doi: 10.1002/bies.950140103.

15. Bradbury EM, Inglis RJ, Matthews HR, Sarner N (1973) Phosphorylation of very-lysine-rich histone in Physarum polycephalum. Correlation with chromosome condensation. Eur J Biochem 33: 131–139

16. Chen JS, Faller DV (2005) Histone deacetylase inhibition-mediated post-translational elevation of p27KIP1 protein levels is required for G_1_ arrest in fibroblasts. J Cell Physiol 202: 87–99

17. Choppakatla P, Dekker B, Cutts EE, Vannini A, Dekker J, Funabiki H (2021) Linker histone H1.8 inhibits chromatin binding of condensins and DNA topoisomerase II to tune chromosome length and individualization. Elife 18:10:e68918. PMID: 34406118. doi: 10.7554/eLife.68918.

18. Chou YH, Rosevear E, Goldman RD (1989) Phosphorylation and disassembly of intermediate filaments in mitotic cells. Proc Natl Acad Sci USA 86: 1885–1889.

19. Chu L, Liang Z, Mukhina M, Fisher J, Vincenten N, Zhang Z, Hutchinson J, Zickler D, Kleckner N (2020) The 3D Topography of Mitotic Chromosomes. Mol Cell 79:902–916.e6. PMID: 32768407 doi: 10.1016/j.molcel.2020.07.002.

20. Coco-Martin JM, Begg AC (1997) Detection of radiation-induced chromosome aberrations using fluorescence in situ hybridization in drug-induced premature chromosome condensations of tumour cell lines with different radiosensitivities. Int J Radiat Biol 71: 265–273

21. Conti D, Esposito Verza A, Pesenti ME, Cmentowski V, Vetter IR, Pan D, Musacchio A (2024) Role of protein kinase PLK1 in the epigenetic maintenance of centromeres. Science 385:1091–1097. PMID: 39236163. doi: 10.1126/science.ado5178.

22. Contreras A, Hale TK, Stenoien DL, Rosen JM, Mancini MA, Herrera RE (2003) The dynamic mobility of histone H1 is regulated by cyclin/CDK phosphorylation. Mol Cell Biol 23: 8626–8636.

23. Cooper S (2003) Rethinking synchronization of mammalian cells for cell cycle analysis. Cell Mol Life Sci 60: 1099–1106

24. Dai J, Sultan S, Taylor SS, Higgins JMG (2005) The kinase haspin is required for mitotic histone H3 Thr 3 phosphorylation and normal metaphase chromosome alignment. Genes Dev 19: 472–488. doi: 10.1101/gad.1267105

25. Darzynkiewicz A, Traganos F, Xue S-B, Melamed MR (1981) Effect of n-butyrate on cell cycle progression and in situ chromatin structure in L1210 cells. Exp Cell Res 136: 279–293

26. Davidson IF, Bauer B, Goetz D, Tang W, Wutz G, Peters J-M (2019) DNA loop extrusion by human cohesion. Science 366:1338–1345. PMID: 31753851. doi: 10.1126/science.aaz3418.

27. de Castro IJ, Gokhan E, Vagnarelli P (2016) Resetting a functional G_1_ nucleus after mitosis. Chromosoma 125: 607–19. doi: 10.1007/s00412-015-0561-6.

28. de Castro IJ, Gil RS, Ligammari L, Di Giacinto ML, Vagnarelli P (2017) CDK1 and PLK1 coordinate the disassembly and reassembly of the nuclear envelope in vertebrate mitosis. Oncotarget 9: 7763–7773. doi: 10.18632/oncotarget.23666

29. Dekker J, Mirny LA (2024) The chromosome folding problem and how cells solve it. Cell 187:6424–6450. PMID: 39547207. doi: 10.1016/j.cell.2024.10.026.

30. Deterding LJ, Bunger MK, Banks GC, Tomer KB, Archer TK (2008) Global changes in and characterization of specific sites of phosphorylation in mouse and human histone H1 isoforms upon CDK inhibitor treatment using mass spectrometry. J Proteome Res 7: 2368–2379

31. Doree M, Galas S (1994) The cyclin-dependent protein kinases and the control of cell division. FASEB J 8: 1114–1121

32. Dupont C, Chahar D, Trullo A, Gostan T, Surcis C, Grimaud C, Fisher D, Feil R, Llères D (2023) Evidence for low nanocompaction of heterochromatin in living embryonic stem cells. EMBO J 42:e110286. doi: 10.15252/embj.2021110286

33. DuPraw EJ (1965) The organization of nuclei and chromosomes in honeybee embryonic cells. Proc Natl Acad Sci USA 53: 161–168. doi: 10.1073/pnas.53.1.161

34. DuPraw EJ (1970) DNA and Chromosomes. Holt, Rinehart and Winston, New York.

35. Earnshaw WC, Rothfield N (1985) Identification of a family of human centromere proteins using autoimmune sera from patients with scleroderma. Chromosoma 91:313–21. PMID: 2579778. doi: 10.1007/BF00328227.

36. Eltsov M, MacLellan KM, Maeshima K, Dubochet J (2008) Analysis of cryo-electron microscopy images does not support the existence of 30-nm chromatin fibers in mitotic chromosomes in situ. Proc Natl Acad Sci USA 105: 19732–19737. 10.1073/pnas.08100571

37. Ferrari S (2006) Protein kinases controlling the onset of mitosis. Cell Mol Life Sci 63: 781–795.

38. Foisner R, Gerace L (1993) Integral membrane proteins of the nuclear envelope interact with lamins and chromosomes, and binding is modulated by mitotic phosphorylation. Cell 73: 1267–1279

39. Forte G, Boteva L, Conforto F, Gilbert N, Cook PR, Marenduzzo D (2024) Bridging condensins mediate compaction of mitotic chromosomes. J Cell Biol 223:e202209113. PMID: 37976091. doi: 10.1083/jcb.202209113.

40. Forte G, Boteva L, Gilbert N, Cook PR, Marenduzzo D (2025) Bridging-mediated compaction of mitotic chromosomes. Nucleus 16:2497765. PMID: 40340634. doi: 10.1080/19491034.2025.2497765

41. Frisa PS, Jacobberger JW (2009) Cell cycle-related cyclin B1 quantification. PLoS One 4: e7064 doi:10.1371/journal.pone.0007064

42. Fung TK, Poon RYC (2005) A roller coaster ride with the mitotic cyclins. Sem Cell Devel Biol 16: 335–342

43. Gadbois DM, Hamaguchi JR, Swank RA, Bradbury EM (1992) Staurosporine is a potent inhibitor of p34^cdc2^ and p34^cdc2^-like kinases. Biochem Biophys Res Commun 184: 80–85

44. Gadea BB, Ruderman JV (2005) Aurora kinase inhibitor ZM447439 blocks chromosome-induced spindle assembly, the completion of chromosome condensation, and the establishment of the spindle integrity checkpoint in Xenopus egg extracts. Mol Biol Cell 16: 1305–1318.

45. Ganji M, Shaltiel IA, Bisht S, Kim E, Kalichava A, Haering CH, Dekker C (2018) Real-time imaging of DNA loop extrusion by condensing. Science 360:102–105. PMID: 29472443. doi: 10.1126/science.aar7831.

46. Garcia BA, Busby SA, Barber CM, Shabanowitz J, Allis CD, Hunt DF (2004) Characterization of phosphorylation sites on histone H1 isoforms by tandem mass spectrometry. J Proteome Res 3: 1219–1227

47. Gautier J, Norbury C, Lohka M, Nurse P, Maller J (1988) Purified maturation-promoting factor contains the product of a Xenopus homolog of the fission yeast cell cycle control gene cdc2+. Cell 54:433–9. PMID: 3293803. doi: 10.1016/0092-8674(88)90206-1.

48. Gautier J, Minshull J, Lohka M, Glotzer M, Hunt T, Maller JL (1990) Cyclin is a component of maturation-promoting factor from Xenopus. Cell 60:487–94. PMID: 1967981. doi: 10.1016/0092-8674(90)90599-a.

49. Gerguri T, Fu X, Kakui Y, Khatri BS, Barrington C, Bates PA, Uhlmann F (2021) Comparison of loop extrusion and diffusion capture as mitotic chromosome formation pathways in fission yeast. Nucleic Acids Res 49:1294–1312. PMID: 33434270. doi: 10.1093/nar/gkaa1270.

50. Gibcus JH, Samejima K, Goloborodko A, Samejima I, Naumova N, Nuebler J, Kanemaki M, Xie L, Paulson JR, Earnshaw WC, Mirny LA, Dekker J (2018) A pathway for mitotic chromosome formation. Science 359: eaao6135. PMID: 29348367. doi: 10.1126/science.aao6135

51. Gibson BA, Doolittle LK, Schneider MWG, Jensen LE, Gamarra N, Henry L, Gerlich DW, Redding S, Rosen MK (2019) Organization of Chromatin by Intrinsic and Regulated Phase Separation. Cell 179:470–484.e21. PMID: 31543265. doi: 10.1016/j.cell.2019.08.037

52. Glass JR, Gerace L (1990) Lamins A and C bind and assemble at the surface of mitotic chromosomes. J Cell Biol 111: 1047–1057

53. Golfier S, Quail T, Kimura H, Brugués J (2020) Cohesin and condensin extrude DNA loops in a cell cycle-dependent manner. Elife 12:9:e53885. PMID: 32396063. doi: 10.7554/eLife.53885.

54. Goloborodko A, Marko JF, Mirny LA (2016) Chromosome Compaction by Active Loop Extrusion. Biophys J 110:2162–8. PMID: 27224481. doi: 10.1016/j.bpj.2016.02.041.

55. Gong J, Traganos F, Darzynkiewicz A (1994) Use of the cyclin E restriction point to map cell arrest in G_1_ induced by n-butyrate, cycloheximide, staurosporine, lovastatin, mimosine and quercetin. Int J Oncol 4: 803–808

56. Gotoh E (2007) Visualizing the dynamics of chromosome structure formation coupled with DNA replication. Chromosoma 116: 453–462

57. Gotoh E (2009) Drug-induced premature chromosome condensation (PCC) protocols: cytogenetic approaches in mitotic chromosome and interphase chromatin. Methods Mol Biol 523: 83–92

58. Gotoh E, Asakawa Y, Kosaka H (1995) Inhibition of protein serine/threonine phosphatases directly induces premature chromosome condensation in mammalian somatic cells. Biomedical Res 16: 63–68

59. Gottesfeld JM, Forbes DJ (1997) Mitotic repression of the transcriptional machinery. Trends Biochem Sci 22: 197–202

60. Gowdy PM, Anderson HJ, Roberge M (1998) Entry into mitosis without Cdc2 kinase activation. J Cell Sci 111: 3401–3410.

61. Gurley LR, D’Anna JA, Barham SS, Deaven LL, Tobey RA (1978) Histone phosphorylation and chromatin structure during mitosis in Chinese hamster cells. Eur J Biochem 84: 1–15. PMID: 206429. doi: 10.1111/j.1432-1033.1978.tb12135.x.

62. Hall LL, Th’ng JPH, Guo XW, Teplitz RL, Bradbury EM (1996) A brief staurosporine treatment of mitotic cells triggers premature exit from mitosis and polyploid cell formation. Cancer Res 56: 3551–3559

63. Han F, Zhang H, Zeng C, Peng A, Xue S-B (1987) Reversibility of the early G_1_ blocking of the asynchronous and M-synchronized HeLa cells by butyrate. Beijing Shifan Daxue Xuebao, Ziran Kexueban 1987: 76–81

64. Harlow E, Lane D (1999) Using Antibodies: A Laboratory Manual, Vol. 286, Cold Spring Harbor Laboratory Press.

65. Heald R, McKeon F (1990) Mutations of phosphorylation sites in lamin A that prevent nuclear lamin disassembly in mitosis. Cell 61: 579–589

66. Hegemann B, Hutchins JRA, Hudecz O, Novatchkova M, Rameseder J, Sykora MM, Liu S, Mazanek M, Lénárt P, Hériché J-K, Poser I, Kraut N, Hyman AA, Yaffe MB, Mechtler K, Peters J-M (2011) Systematic Phosphorylation Analysis of Human Mitotic Protein Complexes. Sci Signal 4: rs12. doi: 10.1126/scisignal.2001993

67. Heintz N, Sive HL, Roeder RG (1983) Regulation of human histone gene expression: kinetics of accumulation and changes in the rate of synthesis and in the half-lives of individual histone mRNAs during the HeLa cell cycle. Mol Cell Biol 3:539–550. doi: 10.1128/mcb.3.4.539-550.1983

68. Hendzel MJ, Wei Y, Mancini MA, Van Hooser A, Ranalli T, Brinkley BR, Bazett-Jones DP, Allis CD (1997) Mitosis-specific phosphorylation of histone H3 initiates primarily within pericentromeric heterochromatin during G2 and spreads in an ordered fashion coincident with mitotic chromosome condensation. Chromosoma 106:348–60. PMID: 9362543. doi: 10.1007/s004120050256.

69. Hirano T (2016) Condensin-Based Chromosome Organization from Bacteria to Vertebrates. Cell 164:847–57. PMID: 26919425. doi: 10.1016/j.cell.2016.01.033.

70. Hochegger H, Dejsuphong D, Sonoda E, Saberi A, Rajendra E, Kirk J, Hunt T, Takeda S (2007) An essential role for Cdk1 in S phase control is revealed via chemical genetics in vertebrate cells. J Cell Biol 178:257–68. PMID: 17635936. doi: 10.1083/jcb.200702034.

71. Hoencamp C, Rowland BD (2023) Genome control by SMC complexes. Nat Rev Mol Cell Biol 24:633–650. PMID: 37231112. doi: 10.1038/s41580-023-00609-8.

72. Holt LJ, Tuch BB, Villén J, Johnson AD, Gygi SP, Morgan DO (2009) Global analysis of Cdk1 substrate phosphorylation sites provides insights into evolution. Science 325:1682–6. PMID: 19779198. doi: 10.1126/science.1172867.

73. Houlard M, Cutts EE, Shamim MS, Godwin J, Weisz D, Aiden AP, Aiden EL, Schermelleh L, Vannini A, Nasmyth K (2021) MCPH1 inhibits Condensin II during interphase by regulating its SMC2-Kleisin interface. Elife 10:e73348. PMID: 34850681. doi: 10.7554/eLife.73348.

74. Hsu J-Y, Sun Z-W, Li X, Reuben M, Tatchell K, Bishop DK, Grushcow JM, Brame CJ, Caldwell JA, Hunt DF, Lin R, Smith MM, Allis CD (2000) Mitotic phosphorylation of histone H3 is governed by Ipl1/aurora kinase and Glc7/PP1 phosphatase in budding yeast and nematodes. Cell 102: 279–291. PMID: 10975519.

75. Huang JN, Park I, Ellingson E, Littlepage LE, Pellman D (2001) Activity of the APC^Cdh1^ form of the anaphase-promoting complex persists until S-phase and prevents the premature expression of Cdc20p. J Cell Biol 154: 85–94

76. Hudson DF, Vagnarelli P, Gassmann R, Earnshaw WC (2003) Condensin is required for nonhistone protein assembly and structural integrity of vertebrate mitotic chromosomes. Devel Cell 5: 323–336

77. Huguet F, Flynn S, Vagnarelli P (2019) The Role of Phosphatases in Nuclear Envelope Disassembly and Reassembly and Their Relevance to Pathologies. Cells 8: 687–699. doi: 10.3390/cells8070687.

78. Ishihara H, Martin BL, Brautigan DL, Karaki H, Ozaki H, Kato Y, Fusetani N, Watabe S, Hashimoto K, Uemura D, Hartshorne DJ (1989) Calyculin A and okadaic acid: inhibitors of protein phosphatase activity. Biochem Biophys Res Commun 159: 871–877

79. Janissen R, Barth R, Davidson IF, Peters JM, Dekker C (2024) All eukaryotic SMC proteins induce a twist of -0.6 at each DNA loop extrusion step. Sci Adv 10: eadt1832. doi: 10.1126/sciadv.adt1832.

80. Jansen LET, Black BE, Foltz DR, Cleveland DW (2007) Propagation of centromeric chromatin requires exit from mitosis. J Cell Biol 176:795–805. PMID: 17339380 doi: 10.1083/jcb.200701066.

81. Johnson RT, Rao PN (1970) Mammalian cell fusion: induction of premature chromosome condensation in interphase nuclei. Nature 226: 717–722

82. Joti Y, Hikima T, Nishino Y, Kamada F, Hihara S, Takata H, Ishikawa T, Maeshima K (2012) Chromosomes without a 30-nm chromatin fiber. Nucleus 3: 404–410. 10.4161/nucl.21222

83. Kanda R, Eguchi-Kasai K, Hayata I (1999) Phosphatase inhibitors and premature chromosome condensation in human peripheral lymphocytes at different cell-cycle phases. Somat Cell Mol Genet 25: 1–8

84. Kaszás E, Cande WZ (2000) Phosphorylation of histone H3 is correlated with changes in the maintenance of sister chromatid cohesion during meiosis in maize, rather than the condensation of the chromatin. J Cell Sci 113:3217–26. PMID: 10954420. doi: 10.1242/jcs.113.18.3217.

85. Kawashima SA, Yamagishi Y, Honda T, Ishiguro K-I, Watanabe Y (2010) Phosphorylation of H2A by Bub1 prevents chromosomal instability through localizing shugoshin. Science 327: 172–177. doi: 10.1126/science.1180189

86. Keaton JM, Workman BG, Xie L, Paulson JR (2023) Exit from Mitosis in Budding Yeast: Protein Phosphatase 1 is Required Downstream from Cdk1 Inactivation. Chromosome Res 31: 27. doi: 10.1007/s10577-023-09736-6

87. Kimura K, Hirano M, Kobayashi R, Hirano T (1998) Phosphorylation and activation of 13S condensin by Cdc2 in vitro. Science 282:487–90. PMID: 9774278. doi: 10.1126/science.282.5388.487.

88. Kimura K, Rybenkov VV, Crisona NJ, Hirano T, Cozzarelli NR (1999) 13S condensin actively reconfigures DNA by introducing global positive writhe: implications for chromosome condensation. Cell 98:239–48. PMID: 10428035. doi: 10.1016/s0092-8674(00)81018-1.

89. Kornberg RD, Lorch Y (1999) Twenty-five years of the nucleosome, fundamental particle of the eukaryotic chromosome. Cell 98: 285–294

90. Labbé JC, Picard A, Peaucellier G, Cavadore JC, Nurse P, Doree M (1989) Purification of MPF from starfish: identification as the H1 histone kinase p34cdc2 and a possible mechanism for its periodic activation. Cell 57: 253–263

91. Lake RS, Salzman NP (1972) Occurrence and properties of a chromatin-associated F1- histone phosphokinase in mitotic Chinese hamster cells. Biochem 11: 4817–4826.

92. Langan TA, Gautier J, Lohka M, Hollingsworth R, Moreno S, Nurse P, Maller J, Sclafani RA (1989) Mammalian growth-associated H1 histone kinase: a homolog of cdc2+/CDC28 protein kinases controlling mitotic entry in yeast and frog cells. Mol Cell Biol 9: 3860–3868

93. Langmore JP, Paulson JR (1983) Low angle X-ray diffraction studies of chromatin structure in vivo and in isolated nuclei and metaphase chromosomes. J Cell Biol 96: 1120–1131.

94. Lau HW, Ma HT, Yeung TK, Tam MY, Zheng D, Chu SK, Poon RYC (2021) Quantitative differences between cyclin-dependent kinases underlie the unique functions of CDK1 in human cells. Cell Reports 37: 109808.

95. Laurell E, Beck K, Krupina K, Theerthagiri G, Bodenmiller B, Horvath P, Aebersold R, Antonin W, Kutay U (2011) Phosphorylation of Nup98 by multiple kinases is crucial for NPC disassembly during mitotic entry. Cell 144: 539–50. doi: 10.1016/j.cell.2011.01.012.

96. Lee J, Chen L-F, Gaudin S, Gupta K, Spakowitz A, Boettiger AN (2025) Kinetic organization of the genome revealed by ultra-resolution, multiscale live imaging. bioRxiv [Preprint]. 2025 Apr 1:2025.03.27.645817. doi: 10.1101/2025.03.27.645817. PMID: 40236138

97. Lindqvist A, Rodriguez-Bravo V, Medema RH (2009) The decision to enter mitosis: feedback and redundancy in the mitotic entry network. J Cell Biol 185: 193–202

98. Lipp JJ, Hirota T, Poser I, Peters J-M (2007) Aurora B controls the association of condensin I but not condensin II with mitotic chromosomes. J Cell Sci 120:1245–55. PMID: 17356064. doi: 10.1242/jcs.03425.

99. Llères D, Swift S, Lamond AI (2007) Detecting protein-protein interactions in vivo with FRET using multiphoton fluorescence lifetime imaging microscopy (FLIM). In Current Protocols in Cytometry, Robinson JP et al. (eds.) Chapter 12: Unit12.10. doi: 10.1002/0471142956.cy1210s42

100. Llères D, James J, Swift S, Norman DG, Lamond AI (2009) Quantitative analysis of chromatin compaction in living cells using FLIM-FRET. J Cell Biol 187:481–96. PMID: 19948497. doi: 10.1083/jcb.200907029

101. Lohka MJ, Hayes MK, Maller JL (1988) Purification of maturation-promoting factor, an intracellular regulator of early mitotic events. Proc Natl Acad Sci U S A 85:3009–13. PMID: 3283736. doi: 10.1073/pnas.85.9.3009.

102. Luger K, Mäder AW, Richmond RK, Sargent DF, Richmond TJ (1997) Crystal structure of the nucleosome core particle at 2.8 Å resolution. Nature 389: 251–260

103. Maeshima K, Hihara S, Eltsov M (2010) Chromatin structure: does the 30-nm fibre exist in vivo? Curr Op Cell Biol 22: 291–297

104. Malik R, Nigg EA, Körner R (2008) Comparative conservation analysis of the human mitotic phosphoproteome. Bioinformatics 24: 1426–1432. 10.1093/bioinformatics/btn197

105. Marsden MP, Laemmli UK (1979) Metaphase chromosome structure: evidence for a radial loop model. Cell 17:849–858. doi: 10.1016/0092-8674(79)90325-8.

106. McCullock TW, MacLean DM, Kammermeier PJ (2020) Comparing the performance of mScarlet-I, mRuby3, and mCherry as FRET acceptors for mNeonGreen. PLoS One 15:e0219886. doi: 10.1371/journal.pone.0219886

107. Medappa KC, McLean C, Rueckert RR (1971) On the structure of rhinovirus 1A. Virology 44: 259–270

108. Meijer L, Borgne A, Mulner O, Chong JP, Blow JJ, Inagaki N, Inagaki M, Delcros JG, Moulinoux JP (1997) Biochemical and cellular effects of roscovitine, a potent and selective inhibitor of the cyclin-dependent kinases cdc2, cdk2 and cdk5. Eur J Biochem 243: 527–536

109. Merrill GF (1998) Cell synchronization. Meth Cell Biol 57: 229–249

110. Mineo C, Murakami Y, Ishimi Y, Hanaoka F, Yamada MA (1986) Isolation and analysis of a mammalian temperature-sensitive mutant defective in G_2_ functions. Exp Cell Res 167: 53–62

111. Miura T, Blakely WF (2011) Optimization of calyculin A-induced premature chromosome condensation assay for chromosome aberration studies. Cytometry 79A: 1016-1022

112. Murnion ME, Adams RR, Callister DM, Allis CD, Earnshaw WC, Swedlow JR (2001) Chromatin-associated protein phosphatase 1 regulates aurora-B and histone H3 phosphorylation. J Biol Chem 276:26656–65. PMID: 11350965. doi: 10.1074/jbc.M102288200.

113. Musio A, Mariani T, Frediani C, Ascoli C, Sbrana I (1997) Atomic force microscope imaging of chromosome structure during G-banding treatments. Genome 40: 127–131. Doi: 10.1139/g97-018

114. Naumova N, Imakaev M, Fudenberg G, Zhan Y, Lajoie BR, Mirny LA, Dekker J (2013) Organization of the mitotic chromosome. Science 342:948–53. PMID: 24200812. doi: 10.1126/science.1236083.

115. Nasmyth K (2001) Disseminating the genome: joining, resolving, and separating sister chromatids during mitosis and meiosis. Annu Rev Genet 35:673–745. PMID: 11700297. doi: 10.1146/annurev.genet.35.102401.091334.

116. Nelson DA, Krucher NA, Ludlow JW (1997) High molecular weight protein phosphatase type 1 dephosphorylates the retinoblastoma protein. J Biol Chem 272: 4528–4535

117. Norbury C, Nurse P (1992) Animal cell cycles and their control. Annu Rev Biochem 61: 441–470

118. Ono T, Losada A, Hirano M, Myers MP, Neuwald AF, Hirano T (2003) Differential contributions of condensin I and condensin II to mitotic chromosome architecture in vertebrate cells. Cell 115:109–21. PMID: 14532007. doi: 10.1016/s0092-8674(03)00724-4.

119. Pan D, Klare K, Petrovic A, Take A, Walstein K, Singh P, Rondelet A, Bird AW, Musacchio A (2017) CDK-regulated dimerization of M18BP1 on a Mis18 hexamer is necessary for CENP-A loading. Elife 6:e23352. PMID: 28059702. doi: 10.7554/eLife.23352.

120. Panyim S, Chalkley R (1969) High resolution acrylamide gel electrophoresis of histones. Arch Biochem Biophys 130: 337 – 346

121. Parashara P, Medina-Pritchard B, Abad MA, Sotelo-Parrilla P, Thamkachy R, Grundei D, Zou J, Spanos C, Kumar CN, Basquin C, Das V, Yan Z, Al-Murtadha AA, Kelly DA, McHugh T, Imhof A, Rappsilber J, Jeyaprakash AA (2024) PLK1-mediated phosphorylation cascade activates Mis18 complex to ensure centromere inheritance. Science 385:1098–1104. PMID: 39236175. doi: 10.1126/science.ado8270.

122. Pathak R, Ramakumar A, Subramanian U, Prasanna PGS (2009) Differential radio- sensitivities of human chromosomes 1 and 2 in one donor in interphase- and metaphase-spreads after 60Co γ-irradiation. BMC Med Phys 9:6 (http://www.biomedcentral.com/1756-66549/9/6)

123. Patzlaff JS, Terrenoire E, Turner BM, Earnshaw WC, Paulson JR (2010) Acetylation of core histones in response to HDAC inhibitors is diminished in mitotic HeLa cells. Exp Cell Res 316: 2123–2135. doi: 10.1016/j.yexcr.2010.05.003

124. Paulson JR (1980) Sulfhydryl reagents prevent dephosphorylation and proteolysis of histones in isolated HeLa metaphase chromosomes. Eur J Biochem 111: 189–197

125. Paulson JR (1982) Isolation of chromosome clusters from metaphase-arrested HeLa cells. Chromosoma 85: 571–581.

126. Paulson JR (2007) Inactivation of Cdk1/cyclin B in metaphase-arrested mouse FT210 cells induces exit from mitosis without chromosome segregation or cytokinesis and allows passage through another cell cycle. Chromosoma 116: 215–225

127. Paulson, J.R. (2024) Inducing exit from mitosis by inactivating Cdk1/cyclin B in metaphase-arrested cells: A tool to study mitotic exit and reestablishment of interphase. In Methods in Molecular Biology, Vol. 2874, V. M. Bolanos-Garcia, ed., Humana Press, pp. 183–198. 10.1007/978-1-0716-4236-8_15

128. Paulson JR, Higley LL (1999) Acid-urea polyacrylamide slab gel electrophoresis of proteins: Preventing distortion of gel wells during preelectrophoresis. Anal

129. Paulson JR, Langmore JP (1983) Biochem 268: 157-159 Low angle x-ray diffraction studies of HeLa metaphase chromosomes: effects of histone phosphorylation and chromosome isolation procedure. J Cell Biol 96:1132–1137. doi: 10.1083/jcb.96.4.1132.

130. Paulson JR, Vander Mause ER (2013) Calyculin A induces prematurely condensed chromosomes without histone H1 phosphorylation in mammalian G_1_-phase cells. Adv Biol Chem 3: 36–43.

131. Paulson JR, Patzlaff JS, Vallis AJ (1996) Evidence that the endogenous histone H1 phosphatase in HeLa mitotic chromosomes is protein phosphatase 1, not protein phosphatase 2A. J Cell Sci 109: 1437–1447

132. Paulson JR, Hudson DF, Cisneros-Soberanis F, Earnshaw WC (2021) Mitotic chromosomes. Semin Cell Dev Biol 117: 7–29. doi: 10.1016/j.semcdb.2021.03.014

133. Peter M, Nakagawa J, Doree M, Labbe JC, Nigg EA (1990) In vitro disassembly of the nuclear lamina and M phase-specific phosphorylation of lamins by cdc2 kinase. Cell 61: 591–602

134. Pradhan B, Barth R, Kim E, Davidson IF, Bauer B, van Laar T, Yang W, Ryu J-K, van der Torre J, Peters J-M, Dekker C (2022) SMC complexes can traverse physical roadblocks bigger than their ring size. Cell Rep 41:111491. PMID: 36261017. doi: 10.1016/j.celrep.2022.111491.

135. Prasanna PG, Escalada ND, Blakely WF (2000) Induction of premature chromosome condensation by a phosphatase inhibitor and a protein kinase in unstimulated human peripheral blood lymphocytes: a simple and rapid technique to study chromosome aberrations using specific whole-chromosome DNA hybridization probes for biological dosimetry. Mutat Res 466: 131–141. doi: 10.1016/s1383-5718(00)00011-5

136. Prevo B, Peterman EJG (2014) Förster resonance energy transfer and kinesin motor proteins. Chem Soc Rev 43:1144–1155. doi: 10.1039/c3cs60292c.

137. Rao PN (1990) The discovery (or rediscovery?) of the phenomenon of premature chromosome condensation. BioEssays 12: 193–197.

138. Rata S, Suarez Peredo Rodriguez MF, Joseph S, Peter N, Iturra FE, Yang F, Madzvamuse A, Ruppert JG, Samejima K, Platani M, Alvarez-Fernandez M, Malumbres M, Earnshaw WC, Novak B, Hochegger H (2018) Two interlinked bistable switches govern mitotic control in mammalian cells. Curr Biol 28:3824–3832.e6. PMID: 30449668.doi: 10.1016/j.cub.2018.09.059

139. Rattner JB, Hamkalo BA (1979) Nucleosome packing in interphase chromatin. J Cell Biol 81: 453–457. doi: 10.1083/jcb.81.2.453

140. R Core Team (2025). R: A language and environment for statistical computing. R Foundation for Statistical Computing, Vienna, Austria. Available online at https://www.R-project.org/.

141. Ricci MA, Manzo C, García-Parajo MF, Lakadamyali M, Cosma MP (2015) Chromatin fibers are formed by heterogeneous groups of nucleosomes in vivo. Cell 160: 1145–1158. doi: 10.1016/j.cell.2015.01.054.

142. Robinson PJ, An W, Routh A, Martino F, Chapman L, Roeder RG, Rhodes D (2008) 30 nm chromatin fibre decompaction requires both H4-K16 acetylation and linker histone eviction. J Mol Biol 381: 816–825

143. Samejima K, Booth DG, Ogawa H, Paulson JR, Xie L, Watson CA, Platani M, Kanemaki MT, Earnshaw WC (2018) Functional analysis after rapid degradation of condensins and 3D-EM reveals chromatin volume is uncoupled from chromosome architecture in mitosis. J Cell Sci 131: jcs210187. PMID: 29361541. doi: 10.1242/jcs.210187

144. Samejima I, Spanos C, Samejima K, Rappsilber J, Kustatscher G, Earnshaw WC (2022) Mapping the invisible chromatin transactions of prophase chromosome remodeling. Mol Cell 82:696–708.e4. PMID: 35090599. doi: 10.1016/j.molcel.2021.12.039

145. Samejima K, Gibcus JH, Abraham S, Cisneros-Soberanis F, Samejima I, Beckett AJ, Pučeková N, Abad MA, Spanos C, Medina-Pritchard B, Paulson JR, Xie L, Jeyaprakash AA, Prior IA, Mirny LA, Dekker J, Goloborodko A, Earnshaw WC (2025) Rules of engagement for condensins and cohesins guide mitotic chromosome formation. Science 388(6743):eadq1709. PMID: 40208986. doi: 10.1126/science.adq1709.

146. Schneider MWG, Gibson BA, Otsuka S, Spicer MFD, Petrovic M, Blaukopf C, Langer CCH, Batty P, Nagaraju T, Doolittle LK, Rosen MK, Gerlich DW (2022) A mitotic chromatin phase transition prevents perforation by microtubules. Nature 609:183–190. PMID: 35922507. doi: 10.1038/s41586-022-05027-y.

147. Shintomi K, Hirano Y (2011) The relative ratio of condensin I to II determines chromosome shapes. Genes Dev 25:1464–9 PMID: 21715560. doi: 10.1101/gad.2060311.

148. Shintomi K, Takahashi TS, Hirano T (2015) Reconstitution of mitotic chromatids with a minimum set of purified factors. Nat Cell Biol 17:1014–23. PMID: 26075356. doi: 10.1038/ncb3187.

149. Shintomi K, Inoue F, Watanabe H, Ohsumi K, Ohsugi M, Hirano T (2017) Mitotic chromosome assembly despite nucleosome depletion in Xenopus egg extracts. Science 356:1284–1287. PMID: 28522692. doi: 10.1126/science.aam9702.

150. Shorter J, Warren G (2002) Golgi architecture and inheritance. Annu Rev Cell Dev Biol 18: 479–420

151. Silva MCC, Bodor DL, Stellfox ME, Martins NMC, Hochegger H, Foltz DR, Jansen LET (2012) Cdk activity couples epigenetic centromere inheritance to cell cycle progression. Dev Cell 22:52–63. PMID: 22169070. doi: 10.1016/j.devcel.2011.10.014.

152. Spiller F, Medina-Pritchard B, Abad MA, Wear MA, Molina O, Earnshaw WC, Jeyaprakash AA (2017) Molecular basis for Cdk1-regulated timing of Mis18 complex assembly and CENP-A deposition. EMBO Rep 18:894–905. PMID: 28377371. doi: 10.15252/embr.201643564.

153. Srebniak MI, Trapp GG, Wawrzkiewicz AK, Kazmierczak W, Wiczkowski AK (2005) The usefulness of calyculin A for cytogenetic prenatal diagnosis. J Histochem Cytochem 53: 391–394

154. St-Pierre J, Douziech M, Bazile F, Pascariu M, Bonneil E, Sauvé V, Ratsima H, D’Amours D (2009) Polo kinase regulates mitotic chromosome condensation by hyperactivation of condensin DNA supercoiling activity. Mol Cell 34: 416–426. doi: 10.1016/j.molcel.2009.04.013

155. Stukenberg PT, Lustig KD, McGarry TJ, King RW, Kuang J, Kirschner MW (1997) Systematic identification of mitotic phosphoproteins. Curr Biol 7: 338–348

156. Suzuki M, Piao CQ, Zhao YL, Hei TK (2001) Karyotype analysis of tumorigenic human bronchial epithelial cells transformed by chrysolite asbestos using chemically induced premature chromosome condensation technique. Int J Mol Med 8: 43–47. doi: 10.3892/ijmm.8.1.43.

157. Tane S, Shintomi K, Kinoshita K, Tsubota Y, Yoshida MM, Nishiyama T, Hirano T (2022) Cell cycle-specific loading of condensin I is regulated by the N-terminal tail of its kleisin subunit. Elife 11:e84694. PMID: 36511239. doi: 10.7554/eLife.84694.

158. Th’ng JPH, Wright PS, Hamaguchi J, Lee MG, Norbury CJ, Nurse P, Bradbury EM (1990) The FT210 cell line is a mouse G_2_ phase mutant with a temperature- sensitive cdc2 gene product. Cell 63: 313–324

159. Th’ng JPH, Guo XW, Swank RA, Crissman HA, Bradbury EM (1994) Inhibition of histone phosphorylation by staurosporine leads to chromosome decondensation. J Biol Chem 269: 9568–9573

160. Thoma F, Koller T, Klug A (1979) Involvement of histone H1 in the organization of the nucleosome and of the salt-dependent superstructures of chromatin. J Cell Biol 83: 403–427

161. Thompson LJ, Bollen M, Fields AP (1997) Identification of protein phosphatase 1 as a mitotic lamin phosphatase. J Biol Chem 272: 29693–29697

162. Trimborn M, Schindler D, Neitzel H, Hirano T (2006) Misregulated chromosome condensation in MCPH1 primary microcephaly is mediated by condensin II. Cell Cycle 5:322–6. PMID: 16434882. doi: 10.4161/cc.5.3.2412.

163. Uhlmann F (2025) A unified model for cohesin function in sisterchromatid cohesion and chromatin loop formation. Mol Cell 85:1058–1071. PMID: 40118039. doi: 10.1016/j.molcel.2025.02.005.

164. Van Hooser A, Goodrich DW, Allis CD, Brinkley BR, Mancini MA (1998) Histone H3 phosphorylation is required for the initiation, but not maintenance, of mammalian chromosome condensation. J Cell Sci 111: 3497–3506

165. Vaziri C, Stice L, Faller DV (1998) Butyrate-induced G_1_ arrest results from p21- independent disruption of retinoblastoma protein-mediated signals. Cell Growth Differ 9: 465–74.

166. Ward GE, Kirschner MW (1990) Identification of cell cycle-regulated phosphorylation sites on nuclear lamin C. Cell 61: 561–577

167. Wei Y, Mizzen CA, Cook RG, Gorovsky MA, Allis CD (1998) Phosphorylation of histone H3 at serine 10 is correlated with chromosome condensation during mitosis and meiosis in Tetrahymena. Proc Natl Acad Sci USA 95: 7480–7484

168. Wei Y, Yu L, Bowen J, Gorovsky MA, Allis CD (1999) Phosphorylation of histone H3 is required for proper chromosome condensation and segregation. Cell 97:99–109. PMID: 10199406. doi: 10.1016/s0092-8674(00)80718-7.

169. Wilkins BJ, Rall NA, Ostwal Y, Kruitwagen T, Hiragami-Hamada K, Winkler M, Barral Y, Fischle W, Neumann H (2014) A cascade of histone modifications induces chromatin condensation in mitosis. Science 343: 77–80. 10.1126/science.1244508

170. Wintersberger E, Mudrak I, Wintersberger U (1983) Butyrate inhibits mouse fibroblasts at a control point in the G_1_-phase. J Cell Biochem 21: 239–247

171. Xeros N (1962) Deoxyriboside control and synchronization of mitosis. Nature 194: 682–683

172. Yamagishi Y, Honda T, Tanno Y, Watanabe Y (2010) Two histone marks establish the inner centromere and chromosome bi-orientation. Science 330: 239–243. doi: 10.1126/science.1194498

173. Yamaguchi T, Goto H, Yokoyama T, Silljé H, Hanisch A, Uldschmid A, Takai Y, Oguri T, Nigg EA, Inagaki M (2005) Phosphorylation by Cdk1 induces Plk1-mediated vimentin phosphorylation during mitosis. J Cell Biol 171: 431–436

174. Yamashita D, Shintomi K, Ono T, Gavvovidis I, Schindler D, Neitzel H, Trimborn M, Hirano T (2011) MCPH1 regulates chromosome condensation and shaping as a composite modulator of condensin II. J Cell Biol 194:841–54. PMID: 21911480. doi: 10.1083/jcb.201106141.

175. Yatskevich S, Rhodes J, Nasmyth K (2019) Organization of Chromosomal DNA by SMC Complexes. Annu Rev Genet 53:445–482. PMID: 31577909. doi: 10.1146/annurev-genet-112618-043633.

176. Yoshida MM, Kinoshita K, Shintomi K, Aizawa Y, Hirano T (2024) Regulation of condensin II by self-suppression and release mechanisms. Mol Biol Cell 35:ar21. PMID: 38088875. doi: 10.1091/mbc.E23-10-0392.

177. Zhang T, Paulson JR, Bakhrebah M, Kim JH, Nowell C, Kalitsis P, Hudson DF (2016) Condensin I and II behaviour in interphase nuclei and cells undergoing premature chromosome condensation. Chromosome Res 24:243–69. PMID: 27008552. doi: 10.1007/s10577-016-9519-7

178. Zhang M, Liang C, Chen Q, Yan H, Xu J, Zhao H, Yuan X, Liu J, Lin S, Lu W, Wang F (2020) Histone H2A phosphorylation recruits topoisomerase IIα to centromeres to safeguard genomic stability. EMBO J 39:e101863. doi: 10.15252/embj.2019101863.

179. Zhiteneva A, Bonfiglio JJ, Makarov A, Colby T, Vagnarelli P, Schirmer EC, Matic I, Earnshaw WC (2017) Mitotic post-translational modifications of histones promote chromatin compaction in vitro. Open Biol 7: 170076. PMID: 28903997. doi: 10.1098/rsob.170076

