## Supplemental Fig S1 Legend and Tables for "Calyculin A Induces Premature Chromosome Condensation and Chromatin Compaction in G_1_-Phase HeLa Cells without Histone H1 Phosphorylation"

Supplementary Materials:

Figure Legend:

Fig. S1. Detailed data from the FLIM experiment. (A) Examples of mNeonGreen fluorescence (upper panels) and lifetime images (lower panels) of cells expressing H2B-mNeonGreen and mScarlet-I-H2B (left panel) or H2B-mNeonGreen (right panel): G1(1), G1-PCC (2) and nocodazole-arrested (3) cells. (B) Normalized distributions of lifetime events for the cells shown in (A): H2B-mNeonGreen and mScarlet-I-H2B (red), H2B-mNeonGreen (green). The center of mass for each curve is indicated with a vertical line. (C) Distributions of lifetimes in the FLIM-FRET experiment, as in Fig. 6 (A), but including the non-fluorescent control (wild-type). The background lifetimes for the non-fluorescent control are either random or close to 0.

### Table S1: Primers for Cloning into pcDNA5 FRT Vector

| | |
|---|---|
| FRT5EcoRV_mNeonGreen_rev | ACTGTGCTGGATATCCTACTTGTAAAGTTCATC |
| FRT5EcoRV_mScarlet-I_rev | ACTGTGCTGGATATCCTACTTGTACAGCTCGTCC |
| FRT5HindIII_kozak_mNeonGreen_fw | GTTTAAACTTAAGCTTGCCACCATGGTTAGCAAAGGAGAAG |
| mNeonGreen_linker_mScarlet-I_fw | GATGAACTTTACAAGGGTGGAGGAGGCTCTGGGGGCGTGAGCAAGGGCG |
| mNeonGreen_fw | GTTAGCAAAGGAGAAG |
| mNeonGreen_rev | CTTGTAAAGTTCATC |
| FRT5HindIIIkozakH2Bhumfwnew | GTTTAAACTTAAGCTTGCCACCATGccagagccagcgaag |
| mNeonGreen_GGG_H2Bhum_rev_new | CTTCTCCTTTGCTAACGCCCCCTCCcttagcgctggtgta |
| H2Bhum_GGG_mScarlet-I_rev_new | cttcgctggctctggGCCCCCTCCCTTGTACAGCTCGTCC |
| H2Bhum_fw_new | ccagagccagcgaag |
| FRT5EcoRV_H2Bhum_rev_new | ACTGTGCTGGATATCCTActtagcgctggtgta |
| mNeonGreen_T2A_mScatlet-I_fw | GATGAACTTTACAAGcttgagggcagaggaagtctgctaacatgcggtgacgtggaggagaatcccggccctgctagc GTGAGCAAGGGCG |
| PbEcoRI_ kozak_mNG_fw | ctcaagcttcgAATTCGCCACCATGGTTAGCAAAGGAGAAG |
| PbBamHI_ mSI_rev | agatccggtggatcCCTACTTGTACAGCTCGTCC |
| PbEcoRI_utr_koz_fw | ctcaagcttcgAATTCGTTTAAACTTAAGCTTGCCACC |
| BamHI_utr_H2Brev | agatccggtggatcCACTGTGCTGGATATCCTActtag |
| BamHI_utr-mSI_rev | agatccggtggatcCACTGTGCTGGATATCCTACTTG |
| BamHI_utr_mNG_rev | agatccggtggatcCACTGTGCTGGATATCCTACTTG |

Table S2: Lifetime Measurements: Raw Data

Cell Lines:

| | |
|---|---|
| NK21 | cells expressing H2B-mNeonGreen/mScarlet-I-H2B |
| NK22 | cells expressing H2B-mNeonGreen/mScarlet-I |
| NK23 | cells expressing H2B-mNeonGreen |
| NK13 | cells expressing mNeonGreen-linker-mScarlet-I |
| HeLa | non-fluorescent wild-type HeLa CDK1as |

Lifetimes are given in nanoseconds.

| Lifetime | Cell line | Condition | Replicate |
|---|---|---|---|
| 2.93 | NK21 | G1 | 1 |
| 2.942 | NK21 | G1 | 1 |
| 2.916 | NK21 | G1 | 1 |
| 2.932 | NK21 | G1 | 1 |
| 2.914 | NK21 | G1 | 1 |
| 2.933 | NK21 | G1 | 1 |
| 2.951 | NK21 | G1 | 1 |
| 2.922 | NK21 | G1 | 1 |
| 2.927 | NK21 | G1 | 1 |
| 2.923 | NK21 | G1 | 1 |
| 2.915 | NK21 | G1 | 1 |
| 2.943 | NK21 | G1 | 1 |
| 2.922 | NK21 | G1 | 1 |
| 2.93 | NK21 | G1 | 1 |
| 2.931 | NK21 | G1 | 1 |
| 2.917 | NK21 | G1 | 1 |
| 2.957 | NK21 | G1 | 1 |
| 2.94 | NK21 | G1 | 1 |
| 2.921 | NK21 | G1 | 1 |
| 2.927 | NK21 | G1 | 1 |
| 2.928 | NK22 | G1 | 1 |
| 2.958 | NK22 | G1 | 1 |
| 2.948 | NK22 | G1 | 1 |
| 2.933 | NK22 | G1 | 1 |
| 2.963 | NK22 | G1 | 1 |
| 2.955 | NK22 | G1 | 1 |
| 2.958 | NK22 | G1 | 1 |
| 2.963 | NK22 | G1 | 1 |

| | | | |
|---|---|---|---|
| 2.954 | NK22 | G1 | 1 |
| 2.961 | NK22 | G1 | 1 |
| 2.925 | NK22 | G1 | 1 |
| 2.939 | NK22 | G1 | 1 |
| 2.936 | NK22 | G1 | 1 |
| 2.957 | NK22 | G1 | 1 |
| 2.939 | NK22 | G1 | 1 |
| 2.95 | NK22 | G1 | 1 |
| 2.943 | NK22 | G1 | 1 |
| 2.961 | NK22 | G1 | 1 |
| 2.959 | NK22 | G1 | 1 |
| 2.942 | NK22 | G1 | 1 |
| 2.94 | NK23 | G1 | 1 |
| 2.94 | NK23 | G1 | 1 |
| 2.928 | NK23 | G1 | 1 |
| 2.961 | NK23 | G1 | 1 |
| 2.967 | NK23 | G1 | 1 |
| 2.917 | NK23 | G1 | 1 |
| 2.948 | NK23 | G1 | 1 |
| 2.952 | NK23 | G1 | 1 |
| 2.943 | NK23 | G1 | 1 |
| 2.929 | NK23 | G1 | 1 |
| 2.959 | NK23 | G1 | 1 |
| 2.94 | NK23 | G1 | 1 |
| 2.965 | NK23 | G1 | 1 |
| 2.942 | NK23 | G1 | 1 |
| 2.921 | NK23 | G1 | 1 |
| 2.962 | NK23 | G1 | 1 |
| 2.95 | NK23 | G1 | 1 |
| 2.962 | NK23 | G1 | 1 |
| 2.947 | NK23 | G1 | 1 |
| 2.959 | NK23 | G1 | 1 |
| 2.752 | NK13 | G1 | 1 |
| 2.786 | NK13 | G1 | 1 |
| 2.753 | NK13 | G1 | 1 |
| 2.772 | NK13 | G1 | 1 |
| 2.782 | NK13 | G1 | 1 |
| 2.785 | NK13 | G1 | 1 |
| 2.778 | NK13 | G1 | 1 |
| 2.786 | NK13 | G1 | 1 |
| 2.775 | NK13 | G1 | 1 |
| 2.79 | NK13 | G1 | 1 |

| | | | |
|---|---|---|---|
| 2.69 | NK13 | G1 | 1 |
| 2.791 | NK13 | G1 | 1 |
| 2.8 | NK13 | G1 | 1 |
| 2.801 | NK13 | G1 | 1 |
| 2.794 | NK13 | G1 | 1 |
| 2.806 | NK13 | G1 | 1 |
| 2.803 | NK13 | G1 | 1 |
| 2.805 | NK13 | G1 | 1 |
| 2.777 | NK13 | G1 | 1 |
| 2.8 | NK13 | G1 | 1 |
| 2.29 | HeLa | G1 | 1 |
| 1.726 | HeLa | G1 | 1 |
| 1.792 | HeLa | G1 | 1 |
| 1.654 | HeLa | G1 | 1 |
| 1.553 | HeLa | G1 | 1 |
| 1.544 | HeLa | G1 | 1 |
| 1.528 | HeLa | G1 | 1 |
| 1.453 | HeLa | G1 | 1 |
| 1.251 | HeLa | G1 | 1 |
| 1.343 | HeLa | G1 | 1 |
| 1.518 | HeLa | G1 | 1 |
| 1.588 | HeLa | G1 | 1 |
| 1.657 | HeLa | G1 | 1 |
| 1.531 | HeLa | G1 | 1 |
| 1.722 | HeLa | G1 | 1 |
| 1.687 | HeLa | G1 | 1 |
| 1.676 | HeLa | G1 | 1 |
| 1.705 | HeLa | G1 | 1 |
| 1.691 | HeLa | G1 | 1 |
| 1.551 | HeLa | G1 | 1 |
| 2.908 | NK21 | G1 | 2 |
| 2.874 | NK21 | G1 | 2 |
| 2.88 | NK21 | G1 | 2 |
| 2.849 | NK21 | G1 | 2 |
| 2.891 | NK21 | G1 | 2 |
| 2.885 | NK21 | G1 | 2 |
| 2.9 | NK21 | G1 | 2 |
| 2.906 | NK21 | G1 | 2 |
| 2.873 | NK21 | G1 | 2 |
| 2.897 | NK21 | G1 | 2 |
| 2.874 | NK21 | G1 | 2 |
| 2.897 | NK21 | G1 | 2 |

| | | | |
|---|---|---|---|
| 2.891 | NK21 | G1 | 2 |
| 2.907 | NK21 | G1 | 2 |
| 2.909 | NK21 | G1 | 2 |
| 2.911 | NK21 | G1 | 2 |
| 2.899 | NK21 | G1 | 2 |
| 2.881 | NK21 | G1 | 2 |
| 2.887 | NK21 | G1 | 2 |
| 2.876 | NK21 | G1 | 2 |
| 2.925 | NK22 | G1 | 2 |
| 2.902 | NK22 | G1 | 2 |
| 2.935 | NK22 | G1 | 2 |
| 2.933 | NK22 | G1 | 2 |
| 2.843 | NK22 | G1 | 2 |
| 2.932 | NK22 | G1 | 2 |
| 2.919 | NK22 | G1 | 2 |
| 2.928 | NK22 | G1 | 2 |
| 2.912 | NK22 | G1 | 2 |
| 2.867 | NK22 | G1 | 2 |
| 2.931 | NK22 | G1 | 2 |
| 2.933 | NK22 | G1 | 2 |
| 2.861 | NK22 | G1 | 2 |
| 2.926 | NK22 | G1 | 2 |
| 2.901 | NK22 | G1 | 2 |
| 2.894 | NK22 | G1 | 2 |
| 2.912 | NK22 | G1 | 2 |
| 2.925 | NK22 | G1 | 2 |
| 2.915 | NK22 | G1 | 2 |
| 2.929 | NK22 | G1 | 2 |
| 2.923 | NK23 | G1 | 2 |
| 2.86 | NK23 | G1 | 2 |
| 2.926 | NK23 | G1 | 2 |
| 2.902 | NK23 | G1 | 2 |
| 2.897 | NK23 | G1 | 2 |
| 2.913 | NK23 | G1 | 2 |
| 2.928 | NK23 | G1 | 2 |
| 2.936 | NK23 | G1 | 2 |
| 2.924 | NK23 | G1 | 2 |
| 2.917 | NK23 | G1 | 2 |
| 2.886 | NK23 | G1 | 2 |
| 2.912 | NK23 | G1 | 2 |
| 2.909 | NK23 | G1 | 2 |
| 2.934 | NK23 | G1 | 2 |

| | | | |
|---|---|---|---|
| 2.935 | NK23 | G1 | 2 |
| 2.869 | NK23 | G1 | 2 |
| 2.919 | NK23 | G1 | 2 |
| 2.887 | NK23 | G1 | 2 |
| 2.927 | NK23 | G1 | 2 |
| 2.927 | NK23 | G1 | 2 |
| 2.669 | NK13 | G1 | 2 |
| 2.679 | NK13 | G1 | 2 |
| 2.69 | NK13 | G1 | 2 |
| 2.684 | NK13 | G1 | 2 |
| 2.681 | NK13 | G1 | 2 |
| 2.671 | NK13 | G1 | 2 |
| 2.642 | NK13 | G1 | 2 |
| 2.679 | NK13 | G1 | 2 |
| 2.674 | NK13 | G1 | 2 |
| 2.646 | NK13 | G1 | 2 |
| 2.631 | NK13 | G1 | 2 |
| 2.697 | NK13 | G1 | 2 |
| 2.698 | NK13 | G1 | 2 |
| 2.691 | NK13 | G1 | 2 |
| 2.692 | NK13 | G1 | 2 |
| 2.699 | NK13 | G1 | 2 |
| 2.691 | NK13 | G1 | 2 |
| 2.698 | NK13 | G1 | 2 |
| 2.682 | NK13 | G1 | 2 |
| 2.552 | NK13 | G1 | 2 |
| 0.122 | HeLa | G1 | 2 |
| 0.126 | HeLa | G1 | 2 |
| 0.144 | HeLa | G1 | 2 |
| 0.09 | HeLa | G1 | 2 |
| 0.121 | HeLa | G1 | 2 |
| 0.119 | HeLa | G1 | 2 |
| 0.109 | HeLa | G1 | 2 |
| 0.114 | HeLa | G1 | 2 |
| 0.117 | HeLa | G1 | 2 |
| 0.123 | HeLa | G1 | 2 |
| 0.243 | HeLa | G1 | 2 |
| 0.365 | HeLa | G1 | 2 |
| 0.296 | HeLa | G1 | 2 |
| 0.513 | HeLa | G1 | 2 |
| 0.588 | HeLa | G1 | 2 |
| 0.292 | HeLa | G1 | 2 |

| | | | |
|---|---|---|---|
| 0.415 | HeLa | G1 | 2 |
| 0.403 | HeLa | G1 | 2 |
| 0.184 | HeLa | G1 | 2 |
| 0.437 | HeLa | G1 | 2 |
| 2.916 | NK21 | G1 | 3 |
| 2.856 | NK21 | G1 | 3 |
| 2.88 | NK21 | G1 | 3 |
| 2.893 | NK21 | G1 | 3 |
| 2.865 | NK21 | G1 | 3 |
| 2.871 | NK21 | G1 | 3 |
| 2.901 | NK21 | G1 | 3 |
| 2.888 | NK21 | G1 | 3 |
| 2.898 | NK21 | G1 | 3 |
| 2.895 | NK21 | G1 | 3 |
| 2.868 | NK21 | G1 | 3 |
| 2.903 | NK21 | G1 | 3 |
| 2.889 | NK21 | G1 | 3 |
| 2.895 | NK21 | G1 | 3 |
| 2.89 | NK21 | G1 | 3 |
| 2.898 | NK21 | G1 | 3 |
| 2.894 | NK21 | G1 | 3 |
| 2.892 | NK21 | G1 | 3 |
| 2.913 | NK21 | G1 | 3 |
| 2.904 | NK21 | G1 | 3 |
| 2.927 | NK22 | G1 | 3 |
| 2.885 | NK22 | G1 | 3 |
| 2.907 | NK22 | G1 | 3 |
| 2.921 | NK22 | G1 | 3 |
| 2.932 | NK22 | G1 | 3 |
| 2.904 | NK22 | G1 | 3 |
| 2.922 | NK22 | G1 | 3 |
| 2.929 | NK22 | G1 | 3 |
| 2.916 | NK22 | G1 | 3 |
| 2.925 | NK22 | G1 | 3 |
| 2.841 | NK22 | G1 | 3 |
| 2.898 | NK22 | G1 | 3 |
| 2.922 | NK22 | G1 | 3 |
| 2.908 | NK22 | G1 | 3 |
| 2.887 | NK22 | G1 | 3 |
| 2.917 | NK22 | G1 | 3 |
| 2.902 | NK22 | G1 | 3 |
| 2.874 | NK22 | G1 | 3 |

| | | | |
|---|---|---|---|
| 2.91 | NK22 | G1 | 3 |
| 2.846 | NK22 | G1 | 3 |
| 2.938 | NK23 | G1 | 3 |
| 2.933 | NK23 | G1 | 3 |
| 2.893 | NK23 | G1 | 3 |
| 2.924 | NK23 | G1 | 3 |
| 2.913 | NK23 | G1 | 3 |
| 2.936 | NK23 | G1 | 3 |
| 2.937 | NK23 | G1 | 3 |
| 2.89 | NK23 | G1 | 3 |
| 2.924 | NK23 | G1 | 3 |
| 2.918 | NK23 | G1 | 3 |
| 2.919 | NK23 | G1 | 3 |
| 2.9 | NK23 | G1 | 3 |
| 2.933 | NK23 | G1 | 3 |
| 2.933 | NK23 | G1 | 3 |
| 2.94 | NK23 | G1 | 3 |
| 2.942 | NK23 | G1 | 3 |
| 2.94 | NK23 | G1 | 3 |
| 2.925 | NK23 | G1 | 3 |
| 2.939 | NK23 | G1 | 3 |
| 2.903 | NK23 | G1 | 3 |
| 2.632 | NK13 | G1 | 3 |
| 2.653 | NK13 | G1 | 3 |
| 2.68 | NK13 | G1 | 3 |
| 2.69 | NK13 | G1 | 3 |
| 2.676 | NK13 | G1 | 3 |
| 2.633 | NK13 | G1 | 3 |
| 2.569 | NK13 | G1 | 3 |
| 2.665 | NK13 | G1 | 3 |
| 2.686 | NK13 | G1 | 3 |
| 2.485 | NK13 | G1 | 3 |
| 2.599 | NK13 | G1 | 3 |
| 2.586 | NK13 | G1 | 3 |
| 2.668 | NK13 | G1 | 3 |
| 2.696 | NK13 | G1 | 3 |
| 2.709 | NK13 | G1 | 3 |
| 2.694 | NK13 | G1 | 3 |
| 2.69 | NK13 | G1 | 3 |
| 2.419 | NK13 | G1 | 3 |
| 2.666 | NK13 | G1 | 3 |
| 2.606 | NK13 | G1 | 3 |

| | | | |
|---|---|---|---|
| 0.103 | HeLa | G1 | 3 |
| 0.127 | HeLa | G1 | 3 |
| 0.126 | HeLa | G1 | 3 |
| 0.153 | HeLa | G1 | 3 |
| 0.19 | HeLa | G1 | 3 |
| 0.257 | HeLa | G1 | 3 |
| 0.315 | HeLa | G1 | 3 |
| 0.33 | HeLa | G1 | 3 |
| 0.357 | HeLa | G1 | 3 |
| 0.544 | HeLa | G1 | 3 |
| 0.852 | HeLa | G1 | 3 |
| 0.273 | HeLa | G1 | 3 |
| 0.181 | HeLa | G1 | 3 |
| 0.166 | HeLa | G1 | 3 |
| 0.156 | HeLa | G1 | 3 |
| 0.152 | HeLa | G1 | 3 |
| 0.158 | HeLa | G1 | 3 |
| 0.143 | HeLa | G1 | 3 |
| 0.369 | HeLa | G1 | 3 |
| 0.251 | HeLa | G1 | 3 |
| 2.89 | NK21 | G1-PCC | 1 |
| 2.906 | NK21 | G1-PCC | 1 |
| 2.903 | NK21 | G1-PCC | 1 |
| 2.909 | NK21 | G1-PCC | 1 |
| 2.917 | NK21 | G1-PCC | 1 |
| 2.916 | NK21 | G1-PCC | 1 |
| 2.926 | NK21 | G1-PCC | 1 |
| 2.91 | NK21 | G1-PCC | 1 |
| 2.961 | NK21 | G1-PCC | 1 |
| 2.909 | NK21 | G1-PCC | 1 |
| 2.925 | NK21 | G1-PCC | 1 |
| 2.902 | NK21 | G1-PCC | 1 |
| 2.916 | NK21 | G1-PCC | 1 |
| 2.914 | NK21 | G1-PCC | 1 |
| 2.876 | NK21 | G1-PCC | 1 |
| 2.936 | NK21 | G1-PCC | 1 |
| 2.909 | NK21 | G1-PCC | 1 |
| 2.877 | NK21 | G1-PCC | 1 |
| 2.921 | NK21 | G1-PCC | 1 |
| 2.909 | NK21 | G1-PCC | 1 |
| 2.906 | NK22 | G1-PCC | 1 |
| 2.915 | NK22 | G1-PCC | 1 |

| | | | |
|---|---|---|---|
| 2.926 | NK22 | G1-PCC | 1 |
| 2.949 | NK22 | G1-PCC | 1 |
| 2.92 | NK22 | G1-PCC | 1 |
| 2.933 | NK22 | G1-PCC | 1 |
| 2.953 | NK22 | G1-PCC | 1 |
| 2.939 | NK22 | G1-PCC | 1 |
| 2.906 | NK22 | G1-PCC | 1 |
| 2.944 | NK22 | G1-PCC | 1 |
| 2.903 | NK22 | G1-PCC | 1 |
| 2.96 | NK22 | G1-PCC | 1 |
| 2.936 | NK22 | G1-PCC | 1 |
| 2.984 | NK22 | G1-PCC | 1 |
| 2.996 | NK22 | G1-PCC | 1 |
| 2.909 | NK22 | G1-PCC | 1 |
| 2.976 | NK22 | G1-PCC | 1 |
| 2.975 | NK22 | G1-PCC | 1 |
| 2.956 | NK22 | G1-PCC | 1 |
| 2.937 | NK22 | G1-PCC | 1 |
| 2.951 | NK23 | G1-PCC | 1 |
| 2.975 | NK23 | G1-PCC | 1 |
| 2.936 | NK23 | G1-PCC | 1 |
| 2.936 | NK23 | G1-PCC | 1 |
| 2.95 | NK23 | G1-PCC | 1 |
| 2.97 | NK23 | G1-PCC | 1 |
| 2.935 | NK23 | G1-PCC | 1 |
| 2.921 | NK23 | G1-PCC | 1 |
| 2.899 | NK23 | G1-PCC | 1 |
| 2.988 | NK23 | G1-PCC | 1 |
| 2.882 | NK23 | G1-PCC | 1 |
| 2.958 | NK23 | G1-PCC | 1 |
| 2.933 | NK23 | G1-PCC | 1 |
| 2.996 | NK23 | G1-PCC | 1 |
| 2.882 | NK23 | G1-PCC | 1 |
| 2.978 | NK23 | G1-PCC | 1 |
| 2.958 | NK23 | G1-PCC | 1 |
| 2.994 | NK23 | G1-PCC | 1 |
| 2.945 | NK23 | G1-PCC | 1 |
| 3.011 | NK23 | G1-PCC | 1 |
| 2.648 | NK13 | G1-PCC | 1 |
| 2.715 | NK13 | G1-PCC | 1 |
| 2.719 | NK13 | G1-PCC | 1 |
| 2.707 | NK13 | G1-PCC | 1 |

| | | | |
|---|---|---|---|
| 2.722 | NK13 | G1-PCC | 1 |
| 2.717 | NK13 | G1-PCC | 1 |
| 2.707 | NK13 | G1-PCC | 1 |
| 2.735 | NK13 | G1-PCC | 1 |
| 2.723 | NK13 | G1-PCC | 1 |
| 2.732 | NK13 | G1-PCC | 1 |
| 2.547 | NK13 | G1-PCC | 1 |
| 2.702 | NK13 | G1-PCC | 1 |
| 2.705 | NK13 | G1-PCC | 1 |
| 2.726 | NK13 | G1-PCC | 1 |
| 2.743 | NK13 | G1-PCC | 1 |
| 2.731 | NK13 | G1-PCC | 1 |
| 2.742 | NK13 | G1-PCC | 1 |
| 2.729 | NK13 | G1-PCC | 1 |
| 2.731 | NK13 | G1-PCC | 1 |
| 2.738 | NK13 | G1-PCC | 1 |
| 1.111 | HeLa | G1-PCC | 1 |
| 0.249 | HeLa | G1-PCC | 1 |
| 0.559 | HeLa | G1-PCC | 1 |
| 1.275 | HeLa | G1-PCC | 1 |
| 1.178 | HeLa | G1-PCC | 1 |
| 1.809 | HeLa | G1-PCC | 1 |
| 2.187 | HeLa | G1-PCC | 1 |
| 0.108 | HeLa | G1-PCC | 1 |
| 0.105 | HeLa | G1-PCC | 1 |
| 1.635 | HeLa | G1-PCC | 1 |
| 1.334 | HeLa | G1-PCC | 1 |
| 0.109 | HeLa | G1-PCC | 1 |
| 1.862 | HeLa | G1-PCC | 1 |
| 1.529 | HeLa | G1-PCC | 1 |
| 0.108 | HeLa | G1-PCC | 1 |
| 0.107 | HeLa | G1-PCC | 1 |
| 1.022 | HeLa | G1-PCC | 1 |
| 1.307 | HeLa | G1-PCC | 1 |
| 1.396 | HeLa | G1-PCC | 1 |
| 1.387 | HeLa | G1-PCC | 1 |
| 2.914 | NK21 | G1-PCC | 2 |
| 2.891 | NK21 | G1-PCC | 2 |
| 2.878 | NK21 | G1-PCC | 2 |
| 2.89 | NK21 | G1-PCC | 2 |
| 2.858 | NK21 | G1-PCC | 2 |
| 2.918 | NK21 | G1-PCC | 2 |

| | | | |
|---|---|---|---|
| 2.809 | NK21 | G1-PCC | 2 |
| 2.878 | NK21 | G1-PCC | 2 |
| 2.735 | NK21 | G1-PCC | 2 |
| 2.892 | NK21 | G1-PCC | 2 |
| 2.806 | NK21 | G1-PCC | 2 |
| 2.912 | NK21 | G1-PCC | 2 |
| 2.889 | NK21 | G1-PCC | 2 |
| 2.933 | NK21 | G1-PCC | 2 |
| 2.927 | NK21 | G1-PCC | 2 |
| 2.896 | NK21 | G1-PCC | 2 |
| 2.895 | NK21 | G1-PCC | 2 |
| 2.921 | NK21 | G1-PCC | 2 |
| 2.924 | NK21 | G1-PCC | 2 |
| 2.932 | NK21 | G1-PCC | 2 |
| 2.959 | NK22 | G1-PCC | 2 |
| 2.96 | NK22 | G1-PCC | 2 |
| 2.889 | NK22 | G1-PCC | 2 |
| 2.795 | NK22 | G1-PCC | 2 |
| 2.941 | NK22 | G1-PCC | 2 |
| 2.974 | NK22 | G1-PCC | 2 |
| 2.942 | NK22 | G1-PCC | 2 |
| 2.967 | NK22 | G1-PCC | 2 |
| 2.958 | NK22 | G1-PCC | 2 |
| 2.956 | NK22 | G1-PCC | 2 |
| 2.957 | NK22 | G1-PCC | 2 |
| 2.951 | NK22 | G1-PCC | 2 |
| 2.958 | NK22 | G1-PCC | 2 |
| 2.985 | NK22 | G1-PCC | 2 |
| 2.974 | NK22 | G1-PCC | 2 |
| 2.923 | NK22 | G1-PCC | 2 |
| 2.954 | NK22 | G1-PCC | 2 |
| 2.965 | NK22 | G1-PCC | 2 |
| 3.005 | NK22 | G1-PCC | 2 |
| 2.973 | NK22 | G1-PCC | 2 |
| 2.975 | NK23 | G1-PCC | 2 |
| 2.932 | NK23 | G1-PCC | 2 |
| 2.949 | NK23 | G1-PCC | 2 |
| 2.878 | NK23 | G1-PCC | 2 |
| 2.898 | NK23 | G1-PCC | 2 |
| 2.99 | NK23 | G1-PCC | 2 |
| 2.994 | NK23 | G1-PCC | 2 |
| 2.966 | NK23 | G1-PCC | 2 |

| | | | |
|---|---|---|---|
| 2.969 | NK23 | G1-PCC | 2 |
| 2.901 | NK23 | G1-PCC | 2 |
| 2.889 | NK23 | G1-PCC | 2 |
| 3.008 | NK23 | G1-PCC | 2 |
| 2.965 | NK23 | G1-PCC | 2 |
| 2.959 | NK23 | G1-PCC | 2 |
| 2.99 | NK23 | G1-PCC | 2 |
| 2.977 | NK23 | G1-PCC | 2 |
| 2.916 | NK23 | G1-PCC | 2 |
| 2.974 | NK23 | G1-PCC | 2 |
| 2.961 | NK23 | G1-PCC | 2 |
| 2.995 | NK23 | G1-PCC | 2 |
| 2.611 | NK13 | G1-PCC | 2 |
| 2.72 | NK13 | G1-PCC | 2 |
| 2.733 | NK13 | G1-PCC | 2 |
| 2.735 | NK13 | G1-PCC | 2 |
| 2.674 | NK13 | G1-PCC | 2 |
| 2.677 | NK13 | G1-PCC | 2 |
| 2.732 | NK13 | G1-PCC | 2 |
| 2.708 | NK13 | G1-PCC | 2 |
| 2.723 | NK13 | G1-PCC | 2 |
| 2.739 | NK13 | G1-PCC | 2 |
| 2.551 | NK13 | G1-PCC | 2 |
| 2.643 | NK13 | G1-PCC | 2 |
| 2.727 | NK13 | G1-PCC | 2 |
| 2.749 | NK13 | G1-PCC | 2 |
| 2.715 | NK13 | G1-PCC | 2 |
| 2.696 | NK13 | G1-PCC | 2 |
| 2.747 | NK13 | G1-PCC | 2 |
| 2.752 | NK13 | G1-PCC | 2 |
| 2.718 | NK13 | G1-PCC | 2 |
| 2.657 | NK13 | G1-PCC | 2 |
| 0.116 | HeLa | G1-PCC | 2 |
| 1.409 | HeLa | G1-PCC | 2 |
| 0.908 | HeLa | G1-PCC | 2 |
| 1.221 | HeLa | G1-PCC | 2 |
| 1.166 | HeLa | G1-PCC | 2 |
| 0.922 | HeLa | G1-PCC | 2 |
| 1.684 | HeLa | G1-PCC | 2 |
| 1.56 | HeLa | G1-PCC | 2 |
| 1.592 | HeLa | G1-PCC | 2 |
| 1.63 | HeLa | G1-PCC | 2 |

| | | | |
|---|---|---|---|
| 1.565 | HeLa | G1-PCC | 2 |
| 1.524 | HeLa | G1-PCC | 2 |
| 0.893 | HeLa | G1-PCC | 2 |
| 0.294 | HeLa | G1-PCC | 2 |
| 0.441 | HeLa | G1-PCC | 2 |
| 1.536 | HeLa | G1-PCC | 2 |
| 1.213 | HeLa | G1-PCC | 2 |
| 0.114 | HeLa | G1-PCC | 2 |
| 1.499 | HeLa | G1-PCC | 2 |
| 0.114 | HeLa | G1-PCC | 2 |
| 2.895 | NK21 | G1-PCC | 3 |
| 2.882 | NK21 | G1-PCC | 3 |
| 2.877 | NK21 | G1-PCC | 3 |
| 2.871 | NK21 | G1-PCC | 3 |
| 2.884 | NK21 | G1-PCC | 3 |
| 2.917 | NK21 | G1-PCC | 3 |
| 2.869 | NK21 | G1-PCC | 3 |
| 2.718 | NK21 | G1-PCC | 3 |
| 2.859 | NK21 | G1-PCC | 3 |
| 2.841 | NK21 | G1-PCC | 3 |
| 2.911 | NK21 | G1-PCC | 3 |
| 2.908 | NK21 | G1-PCC | 3 |
| 2.915 | NK21 | G1-PCC | 3 |
| 2.916 | NK21 | G1-PCC | 3 |
| 2.903 | NK21 | G1-PCC | 3 |
| 2.909 | NK21 | G1-PCC | 3 |
| 2.867 | NK21 | G1-PCC | 3 |
| 2.924 | NK21 | G1-PCC | 3 |
| 2.903 | NK21 | G1-PCC | 3 |
| 2.794 | NK21 | G1-PCC | 3 |
| 2.885 | NK22 | G1-PCC | 3 |
| 2.928 | NK22 | G1-PCC | 3 |
| 2.948 | NK22 | G1-PCC | 3 |
| 2.933 | NK22 | G1-PCC | 3 |
| 2.955 | NK22 | G1-PCC | 3 |
| 2.884 | NK22 | G1-PCC | 3 |
| 2.898 | NK22 | G1-PCC | 3 |
| 2.918 | NK22 | G1-PCC | 3 |
| 2.96 | NK22 | G1-PCC | 3 |
| 2.931 | NK22 | G1-PCC | 3 |
| 2.936 | NK22 | G1-PCC | 3 |
| 2.893 | NK22 | G1-PCC | 3 |

| | | | |
|---|---|---|---|
| 2.933 | NK22 | G1-PCC | 3 |
| 2.903 | NK22 | G1-PCC | 3 |
| 2.935 | NK22 | G1-PCC | 3 |
| 2.956 | NK22 | G1-PCC | 3 |
| 2.877 | NK22 | G1-PCC | 3 |
| 2.903 | NK22 | G1-PCC | 3 |
| 2.978 | NK22 | G1-PCC | 3 |
| 2.947 | NK22 | G1-PCC | 3 |
| 2.901 | NK23 | G1-PCC | 3 |
| 2.97 | NK23 | G1-PCC | 3 |
| 2.967 | NK23 | G1-PCC | 3 |
| 2.942 | NK23 | G1-PCC | 3 |
| 2.977 | NK23 | G1-PCC | 3 |
| 2.939 | NK23 | G1-PCC | 3 |
| 2.912 | NK23 | G1-PCC | 3 |
| 2.937 | NK23 | G1-PCC | 3 |
| 2.924 | NK23 | G1-PCC | 3 |
| 2.883 | NK23 | G1-PCC | 3 |
| 2.895 | NK23 | G1-PCC | 3 |
| 2.935 | NK23 | G1-PCC | 3 |
| 2.974 | NK23 | G1-PCC | 3 |
| 2.943 | NK23 | G1-PCC | 3 |
| 2.956 | NK23 | G1-PCC | 3 |
| 2.938 | NK23 | G1-PCC | 3 |
| 2.973 | NK23 | G1-PCC | 3 |
| 2.969 | NK23 | G1-PCC | 3 |
| 2.966 | NK23 | G1-PCC | 3 |
| 2.927 | NK23 | G1-PCC | 3 |
| 2.7 | NK13 | G1-PCC | 3 |
| 2.523 | NK13 | G1-PCC | 3 |
| 2.604 | NK13 | G1-PCC | 3 |
| 2.63 | NK13 | G1-PCC | 3 |
| 2.614 | NK13 | G1-PCC | 3 |
| 2.716 | NK13 | G1-PCC | 3 |
| 2.698 | NK13 | G1-PCC | 3 |
| 2.674 | NK13 | G1-PCC | 3 |
| 2.731 | NK13 | G1-PCC | 3 |
| 2.739 | NK13 | G1-PCC | 3 |
| 2.72 | NK13 | G1-PCC | 3 |
| 2.731 | NK13 | G1-PCC | 3 |
| 2.725 | NK13 | G1-PCC | 3 |
| 2.646 | NK13 | G1-PCC | 3 |

| | | | |
|---|---|---|---|
| 2.707 | NK13 | G1-PCC | 3 |
| 2.709 | NK13 | G1-PCC | 3 |
| 2.74 | NK13 | G1-PCC | 3 |
| 2.729 | NK13 | G1-PCC | 3 |
| 2.759 | NK13 | G1-PCC | 3 |
| 2.727 | NK13 | G1-PCC | 3 |
| 0.814 | HeLa | G1-PCC | 3 |
| 0.109 | HeLa | G1-PCC | 3 |
| 0.477 | HeLa | G1-PCC | 3 |
| 1.258 | HeLa | G1-PCC | 3 |
| 0.111 | HeLa | G1-PCC | 3 |
| 0.111 | HeLa | G1-PCC | 3 |
| 0.116 | HeLa | G1-PCC | 3 |
| 0.113 | HeLa | G1-PCC | 3 |
| 0.112 | HeLa | G1-PCC | 3 |
| 0.072 | HeLa | G1-PCC | 3 |
| 0.112 | HeLa | G1-PCC | 3 |
| 0.112 | HeLa | G1-PCC | 3 |
| 0.877 | HeLa | G1-PCC | 3 |
| 0.116 | HeLa | G1-PCC | 3 |
| 0.114 | HeLa | G1-PCC | 3 |
| 0.112 | HeLa | G1-PCC | 3 |
| 0.109 | HeLa | G1-PCC | 3 |
| 0.118 | HeLa | G1-PCC | 3 |
| 0.779 | HeLa | G1-PCC | 3 |
| 0.644 | HeLa | G1-PCC | 3 |
| 2.782 | NK21 | noc | 1 |
| 2.856 | NK21 | noc | 1 |
| 2.881 | NK21 | noc | 1 |
| 2.833 | NK21 | noc | 1 |
| 2.86 | NK21 | noc | 1 |
| 2.863 | NK21 | noc | 1 |
| 2.869 | NK21 | noc | 1 |
| 2.852 | NK21 | noc | 1 |
| 2.872 | NK21 | noc | 1 |
| 2.862 | NK21 | noc | 1 |
| 2.793 | NK21 | noc | 1 |
| 2.854 | NK21 | noc | 1 |
| 2.857 | NK21 | noc | 1 |
| 2.879 | NK21 | noc | 1 |
| 2.865 | NK21 | noc | 1 |
| 2.881 | NK21 | noc | 1 |

| | | | |
|---|---|---|---|
| 2.866 | NK21 | noc | 1 |
| 2.883 | NK21 | noc | 1 |
| 2.879 | NK21 | noc | 1 |
| 2.876 | NK21 | noc | 1 |
| 2.854 | NK22 | noc | 1 |
| 2.769 | NK22 | noc | 1 |
| 2.865 | NK22 | noc | 1 |
| 2.897 | NK22 | noc | 1 |
| 2.866 | NK22 | noc | 1 |
| 2.875 | NK22 | noc | 1 |
| 2.908 | NK22 | noc | 1 |
| 2.902 | NK22 | noc | 1 |
| 2.892 | NK22 | noc | 1 |
| 2.88 | NK22 | noc | 1 |
| 2.848 | NK22 | noc | 1 |
| 2.879 | NK22 | noc | 1 |
| 2.863 | NK22 | noc | 1 |
| 2.909 | NK22 | noc | 1 |
| 2.895 | NK22 | noc | 1 |
| 2.877 | NK22 | noc | 1 |
| 2.874 | NK22 | noc | 1 |
| 2.883 | NK22 | noc | 1 |
| 2.883 | NK22 | noc | 1 |
| 2.849 | NK22 | noc | 1 |
| 2.842 | NK23 | noc | 1 |
| 2.905 | NK23 | noc | 1 |
| 2.91 | NK23 | noc | 1 |
| 2.901 | NK23 | noc | 1 |
| 2.898 | NK23 | noc | 1 |
| 2.919 | NK23 | noc | 1 |
| 2.913 | NK23 | noc | 1 |
| 2.907 | NK23 | noc | 1 |
| 2.873 | NK23 | noc | 1 |
| 2.901 | NK23 | noc | 1 |
| 2.884 | NK23 | noc | 1 |
| 2.883 | NK23 | noc | 1 |
| 2.908 | NK23 | noc | 1 |
| 2.906 | NK23 | noc | 1 |
| 2.916 | NK23 | noc | 1 |
| 2.92 | NK23 | noc | 1 |
| 2.916 | NK23 | noc | 1 |
| 2.908 | NK23 | noc | 1 |

| | | | |
|---|---|---|---|
| 2.913 | NK23 | noc | 1 |
| 2.908 | NK23 | noc | 1 |
| 2.612 | NK13 | noc | 1 |
| 2.673 | NK13 | noc | 1 |
| 2.704 | NK13 | noc | 1 |
| 2.691 | NK13 | noc | 1 |
| 2.711 | NK13 | noc | 1 |
| 2.694 | NK13 | noc | 1 |
| 2.689 | NK13 | noc | 1 |
| 2.697 | NK13 | noc | 1 |
| 2.699 | NK13 | noc | 1 |
| 2.709 | NK13 | noc | 1 |
| 2.691 | NK13 | noc | 1 |
| 2.639 | NK13 | noc | 1 |
| 2.638 | NK13 | noc | 1 |
| 2.702 | NK13 | noc | 1 |
| 2.71 | NK13 | noc | 1 |
| 2.661 | NK13 | noc | 1 |
| 2.683 | NK13 | noc | 1 |
| 2.683 | NK13 | noc | 1 |
| 2.71 | NK13 | noc | 1 |
| 2.706 | NK13 | noc | 1 |
| 0.393 | HeLa | noc | 1 |
| 1.109 | HeLa | noc | 1 |
| 1.676 | HeLa | noc | 1 |
| 1.727 | HeLa | noc | 1 |
| 1.715 | HeLa | noc | 1 |
| 1.416 | HeLa | noc | 1 |
| 1.329 | HeLa | noc | 1 |
| 2.14 | HeLa | noc | 1 |
| 2.139 | HeLa | noc | 1 |
| 1.884 | HeLa | noc | 1 |
| 3.27 | HeLa | noc | 1 |
| 1.298 | HeLa | noc | 1 |
| 0.802 | HeLa | noc | 1 |
| 0.103 | HeLa | noc | 1 |
| 0.107 | HeLa | noc | 1 |
| 0.111 | HeLa | noc | 1 |
| 0.962 | HeLa | noc | 1 |
| 2.265 | HeLa | noc | 1 |
| 1.633 | HeLa | noc | 1 |
| 2.07 | HeLa | noc | 1 |

| | | | |
|---|---|---|---|
| 2.839 | NK21 | noc | 2 |
| 2.869 | NK21 | noc | 2 |
| 2.835 | NK21 | noc | 2 |
| 2.869 | NK21 | noc | 2 |
| 2.835 | NK21 | noc | 2 |
| 2.881 | NK21 | noc | 2 |
| 2.832 | NK21 | noc | 2 |
| 2.847 | NK21 | noc | 2 |
| 2.806 | NK21 | noc | 2 |
| 2.858 | NK21 | noc | 2 |
| 2.826 | NK21 | noc | 2 |
| 2.884 | NK21 | noc | 2 |
| 2.838 | NK21 | noc | 2 |
| 2.878 | NK21 | noc | 2 |
| 2.821 | NK21 | noc | 2 |
| 2.886 | NK21 | noc | 2 |
| 2.87 | NK21 | noc | 2 |
| 2.853 | NK21 | noc | 2 |
| 2.887 | NK21 | noc | 2 |
| 2.847 | NK21 | noc | 2 |
| 2.843 | NK22 | noc | 2 |
| 2.89 | NK22 | noc | 2 |
| 2.877 | NK22 | noc | 2 |
| 2.914 | NK22 | noc | 2 |
| 2.872 | NK22 | noc | 2 |
| 2.848 | NK22 | noc | 2 |
| 2.902 | NK22 | noc | 2 |
| 2.874 | NK22 | noc | 2 |
| 2.869 | NK22 | noc | 2 |
| 2.873 | NK22 | noc | 2 |
| 2.881 | NK22 | noc | 2 |
| 2.904 | NK22 | noc | 2 |
| 2.873 | NK22 | noc | 2 |
| 2.876 | NK22 | noc | 2 |
| 2.889 | NK22 | noc | 2 |
| 2.902 | NK22 | noc | 2 |
| 2.898 | NK22 | noc | 2 |
| 2.894 | NK22 | noc | 2 |
| 2.892 | NK22 | noc | 2 |
| 2.901 | NK22 | noc | 2 |
| 2.923 | NK23 | noc | 2 |
| 2.916 | NK23 | noc | 2 |

| | | | |
|---|---|---|---|
| 2.845 | NK23 | noc | 2 |
| 2.924 | NK23 | noc | 2 |
| 2.882 | NK23 | noc | 2 |
| 2.916 | NK23 | noc | 2 |
| 2.916 | NK23 | noc | 2 |
| 2.905 | NK23 | noc | 2 |
| 2.92 | NK23 | noc | 2 |
| 2.899 | NK23 | noc | 2 |
| 2.899 | NK23 | noc | 2 |
| 2.906 | NK23 | noc | 2 |
| 2.89 | NK23 | noc | 2 |
| 2.892 | NK23 | noc | 2 |
| 2.925 | NK23 | noc | 2 |
| 2.907 | NK23 | noc | 2 |
| 2.891 | NK23 | noc | 2 |
| 2.907 | NK23 | noc | 2 |
| 2.928 | NK23 | noc | 2 |
| 2.908 | NK23 | noc | 2 |
| 2.68 | NK13 | noc | 2 |
| 2.673 | NK13 | noc | 2 |
| 2.702 | NK13 | noc | 2 |
| 2.695 | NK13 | noc | 2 |
| 2.702 | NK13 | noc | 2 |
| 2.695 | NK13 | noc | 2 |
| 2.678 | NK13 | noc | 2 |
| 2.718 | NK13 | noc | 2 |
| 2.681 | NK13 | noc | 2 |
| 2.712 | NK13 | noc | 2 |
| 2.636 | NK13 | noc | 2 |
| 2.704 | NK13 | noc | 2 |
| 2.707 | NK13 | noc | 2 |
| 2.717 | NK13 | noc | 2 |
| 2.706 | NK13 | noc | 2 |
| 2.702 | NK13 | noc | 2 |
| 2.707 | NK13 | noc | 2 |
| 2.738 | NK13 | noc | 2 |
| 2.713 | NK13 | noc | 2 |
| 2.707 | NK13 | noc | 2 |
| 1.817 | HeLa | noc | 2 |
| 0.904 | HeLa | noc | 2 |
| 1.08 | HeLa | noc | 2 |
| 1.139 | HeLa | noc | 2 |

| | | | |
|---|---|---|---|
| 1.177 | HeLa | noc | 2 |
| 0.727 | HeLa | noc | 2 |
| 1.332 | HeLa | noc | 2 |
| 0.949 | HeLa | noc | 2 |
| 1.266 | HeLa | noc | 2 |
| 1.079 | HeLa | noc | 2 |
| 1.449 | HeLa | noc | 2 |
| 1.309 | HeLa | noc | 2 |
| 0.998 | HeLa | noc | 2 |
| 0.978 | HeLa | noc | 2 |
| 0.94 | HeLa | noc | 2 |
| 0.879 | HeLa | noc | 2 |
| 1.096 | HeLa | noc | 2 |
| 0.751 | HeLa | noc | 2 |
| 1.036 | HeLa | noc | 2 |
| 0.904 | HeLa | noc | 2 |
| 2.829 | NK21 | noc | 3 |
| 2.84 | NK21 | noc | 3 |
| 2.819 | NK21 | noc | 3 |
| 2.853 | NK21 | noc | 3 |
| 2.842 | NK21 | noc | 3 |
| 2.865 | NK21 | noc | 3 |
| 2.872 | NK21 | noc | 3 |
| 2.837 | NK21 | noc | 3 |
| 2.863 | NK21 | noc | 3 |
| 2.823 | NK21 | noc | 3 |
| 2.86 | NK21 | noc | 3 |
| 2.834 | NK21 | noc | 3 |
| 2.881 | NK21 | noc | 3 |
| 2.868 | NK21 | noc | 3 |
| 2.874 | NK21 | noc | 3 |
| 2.876 | NK21 | noc | 3 |
| 2.879 | NK21 | noc | 3 |
| 2.877 | NK21 | noc | 3 |
| 2.877 | NK21 | noc | 3 |
| 2.837 | NK21 | noc | 3 |
| 2.879 | NK22 | noc | 3 |
| 2.897 | NK22 | noc | 3 |
| 2.9 | NK22 | noc | 3 |
| 2.908 | NK22 | noc | 3 |
| 2.882 | NK22 | noc | 3 |
| 2.896 | NK22 | noc | 3 |

| | | | |
|---|---|---|---|
| 2.887 | NK22 | noc | 3 |
| 2.878 | NK22 | noc | 3 |
| 2.864 | NK22 | noc | 3 |
| 2.904 | NK22 | noc | 3 |
| 2.857 | NK22 | noc | 3 |
| 2.889 | NK22 | noc | 3 |
| 2.896 | NK22 | noc | 3 |
| 2.882 | NK22 | noc | 3 |
| 2.863 | NK22 | noc | 3 |
| 2.864 | NK22 | noc | 3 |
| 2.9 | NK22 | noc | 3 |
| 2.894 | NK22 | noc | 3 |
| 2.9 | NK22 | noc | 3 |
| 2.871 | NK22 | noc | 3 |
| 2.905 | NK23 | noc | 3 |
| 2.899 | NK23 | noc | 3 |
| 2.917 | NK23 | noc | 3 |
| 2.897 | NK23 | noc | 3 |
| 2.91 | NK23 | noc | 3 |
| 2.918 | NK23 | noc | 3 |
| 2.913 | NK23 | noc | 3 |
| 2.88 | NK23 | noc | 3 |
| 2.895 | NK23 | noc | 3 |
| 2.915 | NK23 | noc | 3 |
| 2.911 | NK23 | noc | 3 |
| 2.917 | NK23 | noc | 3 |
| 2.907 | NK23 | noc | 3 |
| 2.862 | NK23 | noc | 3 |
| 2.898 | NK23 | noc | 3 |
| 2.923 | NK23 | noc | 3 |
| 2.916 | NK23 | noc | 3 |
| 2.913 | NK23 | noc | 3 |
| 2.896 | NK23 | noc | 3 |
| 2.883 | NK23 | noc | 3 |
| 2.516 | NK13 | noc | 3 |
| 2.656 | NK13 | noc | 3 |
| 2.67 | NK13 | noc | 3 |
| 2.612 | NK13 | noc | 3 |
| 2.663 | NK13 | noc | 3 |
| 2.695 | NK13 | noc | 3 |
| 2.679 | NK13 | noc | 3 |
| 2.665 | NK13 | noc | 3 |

| | | | |
|---|---|---|---|
| 2.673 | NK13 | noc | 3 |
| 2.709 | NK13 | noc | 3 |
| 2.662 | NK13 | noc | 3 |
| 2.662 | NK13 | noc | 3 |
| 2.685 | NK13 | noc | 3 |
| 2.677 | NK13 | noc | 3 |
| 2.655 | NK13 | noc | 3 |
| 2.682 | NK13 | noc | 3 |
| 2.681 | NK13 | noc | 3 |
| 2.618 | NK13 | noc | 3 |
| 2.675 | NK13 | noc | 3 |
| 2.694 | NK13 | noc | 3 |
| 1.989 | HeLa | noc | 3 |
| 0.108 | HeLa | noc | 3 |
| 1.699 | HeLa | noc | 3 |
| 1.722 | HeLa | noc | 3 |
| 0.103 | HeLa | noc | 3 |
| 1.363 | HeLa | noc | 3 |
| 1.495 | HeLa | noc | 3 |
| 1.286 | HeLa | noc | 3 |
| 1.39 | HeLa | noc | 3 |
| 1.603 | HeLa | noc | 3 |
| 1.198 | HeLa | noc | 3 |
| 1.199 | HeLa | noc | 3 |
| 0.967 | HeLa | noc | 3 |
| 0.97 | HeLa | noc | 3 |
| 1.055 | HeLa | noc | 3 |
| 0.951 | HeLa | noc | 3 |
| 0.91 | HeLa | noc | 3 |
| 1.154 | HeLa | noc | 3 |
| 0.885 | HeLa | noc | 3 |
| 0.87 | HeLa | noc | 3 |
