## Supplementary figures and images for "Calyculin A Induces Premature Chromosome Condensation and Chromatin Compaction in G_1_-Phase HeLa Cells without Histone H1 Phosphorylation"

### Supplementary Figure S1

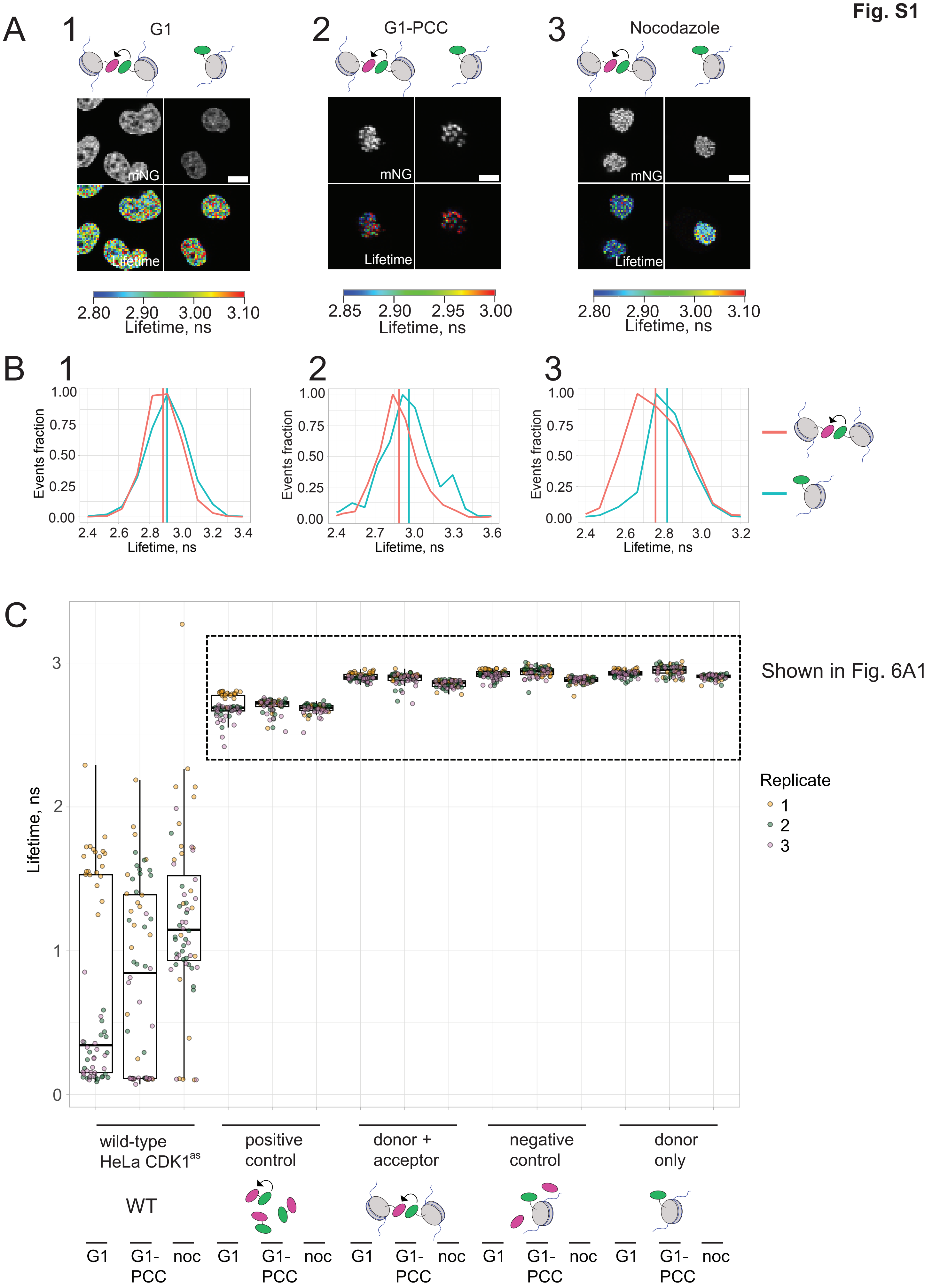
